# Transferrin receptor (Tfr1) ablation in satellite cells impacts skeletal muscle regeneration through the activation of ferroptosis

**DOI:** 10.1101/2020.10.02.323469

**Authors:** Hongrong Ding, Shujie Chen, Xiaohan Pan, Xiaoshuang Dai, Guihua Pan, Ze Li, Xudong Mai, Ye Tian, Susu Zhang, Bingdong Liu, Guangchao Cao, Zhicheng Yao, Xiangping Yao, Liang Gao, Li Yang, Xiaoyan Chen, Jia Sun, Hong Chen, Mulan Han, Yulong Yin, Guohuan Xu, Huijun Li, Weidong Wu, Zheng Chen, Jingchao Lin, Liping Xiang, Yan Lu, Xiao Zhu, Liwei Xie

## Abstract

Satellite cells (SCs) are critical to the postnatal development and skeletal muscle regeneration. Inactivation of SCs is linked with the skeletal muscle loss. Leveraging on the RNAseq screening, transferrin receptor (Tfr1) is identified to be associated with muscle/SC ageing and the declined regeneration potential. Muscle-specific deletion of *Tfr1* results in the growth retardation, metabolic disorder and lethality, shedding light on the importance of Tfr1 in skeletal muscle physiology. Here, our investigation reported that conditional SC-ablation of *Tfr1* leads to the SCs inactivation and skeletal muscle regeneration defects, followed by the labile iron accumulation, *de novo* lipogenesis via fibroadipogenic progenitors (FAPs) and Gpx4/Nrf2-mediated ROS-scavenger defects. These abnormal phenomena, such as Hmox1-mediated myoglobin degradation, Tfr1-Slc39a14 functional switch and the activation of unsaturated fatty acid biosynthesis pathway are orchestrated with the occurrence of ferroptosis in skeletal muscle. Ferroptosis may further prevent SC proliferation and skeletal muscle regeneration. Ferrostatin-1, a ferroptosis inhibitor could not rescue Tfr1-ablation induced ferroptosis. However, intramuscular administration of lentivirus expressing Tfr1 could partially reduce labile iron accumulation, decrease *de novo* lipogenesis and promote skeletal muscle regeneration. Most importantly, Tfr1/Slc39a14 functional switch, labile iron accumulation and fatty acid biosynthesis are recapitulated in aged skeletal muscle of rodents, indicating that ferroptosis occurs in the skeletal muscles of aged rodents. The present study also bridges the gap between pathogenesis of iron and functional defects in the skeletal muscle, providing mechanistic information to develop anti-aging strategies.

**One Sentence Summary:** Conditional ablation of *Tfr1* in satellite cells (SCs) results in the SC inactivation, skeletal muscle regeneration defects, labile iron accumulation, and unsaturated fatty acid biosynthesis, leading to the activation of ferroptosis, which is recapitulated in skeletal muscles of aged rodents to be a new cell death form identified in skeletal muscle and sheds light on the development of novel anti-ageing strategies.

## Introduction

Programmed cell death (PCD), such as apoptosis, autophagy and the newly discovered programmed necrosis, called necroptosis, is a critical process to remove dead, unnecessary or excess cells during organism and tissue development/regeneration*(1)*. Skeletal muscle is a major organ not only supporting movement but also regulating systemic metabolism. Muscle cell death occurs in multiple forms, e.g., apoptosis, necrosis and autophagy*(2)*. Under regenerative conditions, cell death, clearance and regeneration are precisely regulated, while dysregulation of these processes leads to muscular dystrophy, sarcopenia, and pathogenesis in skeletal muscle*(2)*. Necrosis of skeletal muscle occurs under various pathogenic conditions, such as muscular dystrophy and ischemia. However, acute or physiological injuries activate apoptosis, which is regulated by several crucial molecules, such as an anti-apoptotic oncoproteins Bcl2, Caspase 3 and the death receptor Fas/Apo1/Cd95*(3)*. This process is accompanied by the infiltration of inflammatory leukocytes, especially macrophages. At the initial stage of regeneration, M1 macrophages are indispensable for cytokine secretion, fiber debris clearance, iron recycling as well as myoblasts, fibroadipogenic precursor cells (FAPs) and immune cells balancing during skeletal muscle regeneration*(2, 4)*. Upon completion of fiber debris clearance, M1 macrophages are transformed into M2 macrophages contributing to the secretion of anti-inflammatory cytokines and promoting regeneration*(2)*.

Along with these, ferroptosis, a newly identified distinct cell death pathway, is involved in the development of various diseases, such as cancers, ischemia/reperfusion-induced cardiomyopathy, degenerative diseases and stroke. Ferroptosis is an iron- and reactive oxygen species (ROS)-dependent oxidative damage via the accumulation of lipid peroxides*(5, 6)*. Cells experiencing the ferroptosis are characterized by a variety of cytological changes and abnormalities of the mitochondria, including decreased or vanished mitochondrial cristae, a ruptured outer mitochondrial membrane, or a condensed mitochondrial membrane*(7, 8)*. A recent study suggested that during the development of cardiomyopathy death, Nrf2-mediated upregulation of *Hmox1* contributes to free iron release from heme degradation, leading to the lipid peroxidation on mitochondrial lipid bilayer as the major mechanism in ferroptosis-induced heart damage*(9)*. Cardiomyopathy associated ferroptosis is also regulated by *Fth1*-ablation-induced labile iron accumulation and lipid peroxidation, especially under high-iron diet feeding*(10)*. Other diseases, such as liver fibrosis or cirrhosis may be associated with iron-dependent cell death. Patients with liver cirrhosis are diagnosed with higher iron content and lipid peroxidation but lower levels of transferrin (Trf)*(11)*. Liver-specific *Trf*-deletion mice may be susceptible to the development ferroptosis-induced liver fibrosis under a high-iron diet*(11)*. Moreover, the pathogenic property of ferroptosis has not been reported in other tissues, such as skeletal muscle.

Skeletal muscle growth and regeneration rely on a subtype of adult stem cells developed from the mesodermal layer, also called satellite cells (SCs)*(12)*. SCs are considered as adult stem cells because they maintain the self-renewal and remarkable postnatal regenerative potential of skeletal muscle. Quiescent SCs are located between the basal lamina and the plasma lemma of myofibers and are activated by external stimulation or muscle injury, followed by entering the cell cycle, proliferation and differentiation to repair the injured myotubes. This process is regulated by a group of myogenic regulatory factors (MRFs), including but not limited to MyoD and myogenin*(13)*. Some other newly identified transcription factors are also involved in regulating SC physiology and are further extended to the skeletal muscle development and regeneration, such as TTP*(14)*, HIF1/2α*(15, 16)*, and Trp53*(17)*. Dysregulation of these regulatory factors leads to dysfunction of SC and further causes growth and regeneration impairment.

Meanwhile, skeletal muscle growth and regeneration are accompanied by a precise regulation of various nutrients, such amino acids, carbohydrates and minerals. Trace minerals as key nutritional components, play an important role in skeletal muscle physiological function and energy metabolism. Iron, as an essential trace mineral, is required to maintain the appropriate function of skeletal muscles, such as muscle cell differentiation, skeletal muscle growth, and myoglobin synthesis, etc. Iron is released from food and absorbed by epithelial cells of the small intestine. In the form of transferrin bound iron (TBI), it is recognized, internalized and absorbed by the action of transferrin receptor 1 (Tfr1), which is ubiquitously expressed in peripheral tissues such as liver, adipose tissue and skeletal muscles. Thus, Tfr1-mediated iron homeostasis is a rate-limiting step in regulating tissue growth and regeneration. It has been demonstrated that skeletal muscle-specific knockout of *Tfr1* blocks iron absorption and leads to dramatic change in skeletal muscle, liver and adipose tissue metabolism*(18)*. Other studies indicated that skeletal muscle iron is locally recycled with the involvement of myoblasts and macrophages at the different stages of skeletal muscle regeneration. Dysregulation of skeletal muscle iron metabolism, especially labile iron accumulation may impair muscle regeneration *(19)*. Free iron-injection into TA muscle to mimic iron accumulation was shown to induce apoptosis and regeneration defects*(20, 21)*. ROS generated from free iron lead to dysfunction of skeletal muscle. However, the biological function of Tfr1 in SC physiology during skeletal muscle regeneration still remains unknown.

In the present study, leveraging on the RNAseq-screening of gene expression in four different skeletal muscles (TA, EDL, Sol and Gast) at different ages, we identified that *Tfr1* expression is gradually declined with ageing in both skeletal muscle and SCs. SC-specific deletion of *Tfr1* leads to decreased regeneration potential of skeletal muscle, with the phenotype of accumulation of iron and adipocytes, infiltration of macrophages, reduced running ability, mitochondrial stress and metabolic dysfunction. Gene expression profiling of TA muscle from *SC/KO Tfr1^SC/KO^* mice identifies a group of genes associated with ferroptosis, which may cause the irreversible death of a proportion of SCs, possibly due to upregulation of Slc39a14, a nontransferrin bound iron (NTBI) transporter to exacerbate iron-mediated cell death. Unfortunately, Ferrostatin-1, a ferroptosis inhibitor, could not rescue the *Tfr1*-deletion induced ferroptosis, unless Tfr1 function was restored by lentivirus. This genetic model also recapitulates one of the unrecognized ageing-related cell deaths in skeletal muscle with decreased membrane-bound Tfr1 protein but with membrane enrichment of Slc39a14 to uptake NTBI in skeletal muscle of older mice. Taken together, the current investigation identified a previously unrecognized function of the Tfr1-Slc39a14-iron axis in SC during skeletal muscle regeneration and ageing, which sheds light on the development of novel anti-ageing strategies.

## Results

### *Tfr1* expression negatively correlates with skeletal muscle ageing

Skeletal muscle development, growth and maintaining are precisely regulated physiological processes, without which result in sarcopenia*(22)*. To precisely understand these processes, skeletal muscles (TA, EDL, Sol and Gast) were collected from *C57BL/6J* mice across five different ages (Figure S1A). RNAseq followed by bioinformatic analysis identified ~5000 differentially expressed genes (DEGs) among four different muscles (TA:5517 genes; EDL: 4583 genes Sol:5529 genes; Gastr: 5865 genes) between the young (2wk-old) and aged (80wk-old) groups. By plotting the expression of these genes across five ages, we identified a clear trend of gene expression pattern, which was divided into two clusters, a gradually increased (Cluster I) and decreased (Cluster II) expression of genes (Figure S1B-S1E). Of these DEGs of four types of muscles, 2445 DEGs were identified, with 1155 upregulated and 1290 downregulated genes (Figure 1A). As shown by functional analysis of the gene ontology against the biological process gene set, stem cell proliferation and muscle cell differentiation were downregulated in TA muscle of the aged group (Figure 1B). More specifically, iron metabolism-related biological function was declined in the aged group, which was also demonstrated by Gene Set Enrichment Analysis (GSEA) (Figure 1B-C). We also profiled the expression of transition metal ion homeostasis-related genes such as iron, copper and zinc. It was obvious that cellular iron homeostasis-related genes were potentially differentially regulated, while genes involved in copper and zinc ion homeostasis were partially or less differentially regulated across five different ages (Figure S2A-B). Among these genes, *Tfr1* expression was gradually decreased with ageing (from 2-wk to 8-wk, 30-wk, 60-wk and even to 80-wk old) (Figure 1D). Tfr1 is a membrane-bound protein that recognizes transferrin-bound iron (TBI) and is responsible for TBI internalizing in peripheral tissue *(23)*. qPCR and western blotting also confirmed that *Tfr1* mRNA and protein expression was decreased in four different muscles of 8-wk old mice of four different muscle compared to 2-wk old mice, correspondingly accompanied by decreased non-heme iron (Figure S2C). These data indicate that Tfr1-mediated iron absorption is a rate-limiting step in skeletal muscles and may be associated with age-related muscle physiology and function.

**Figure 1.**
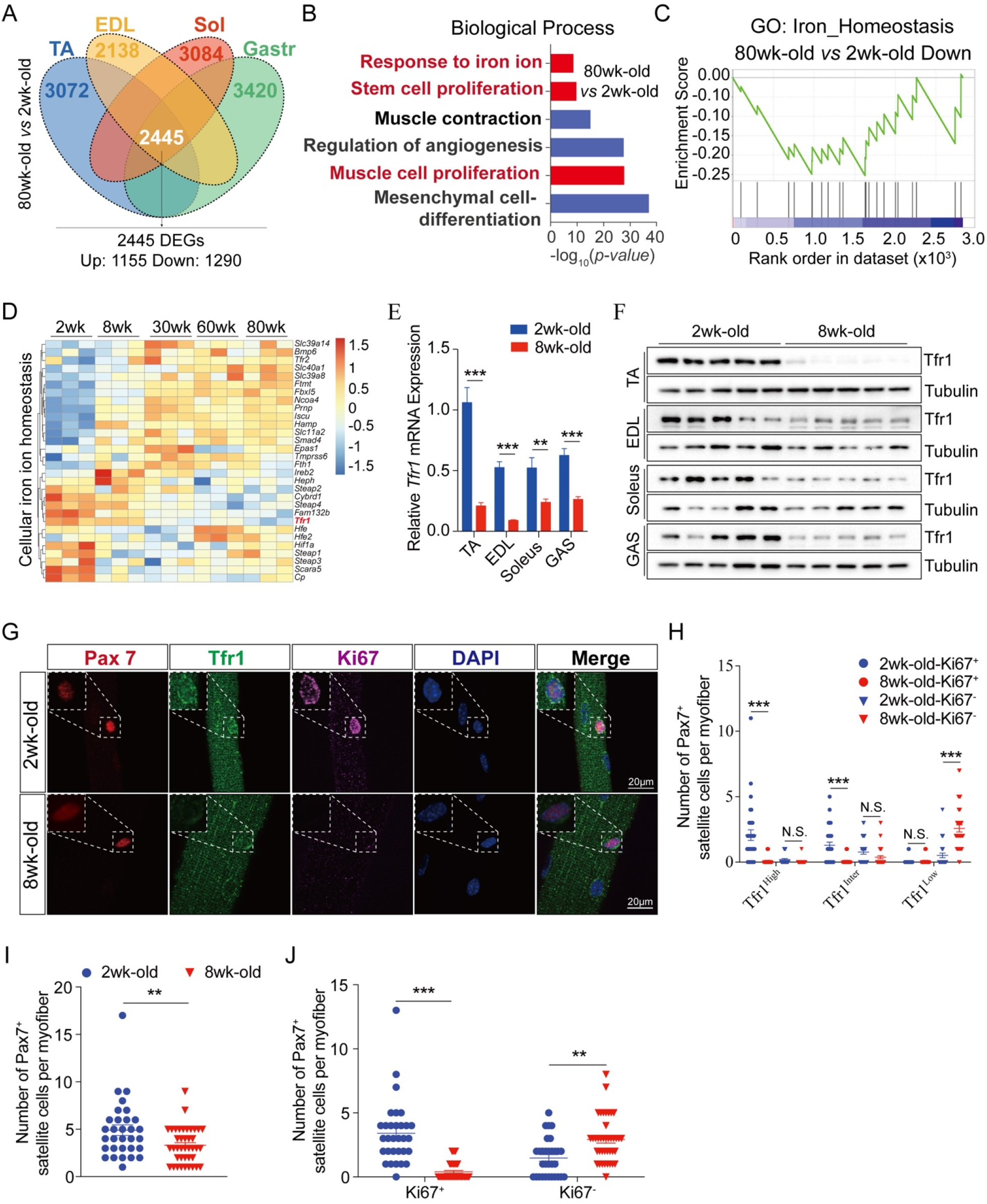
Identification of Tfr1 as a key biomarker regarding skeletal muscle ageing and satellite cells activity. (A) Venn diagraph showing overlapped genes between young (2wk-old) and aged (80wk-old) mice among four types of muscle (TA, EDL, Sol and Gas) (n=3/group); (B) Gene ontology (GO: Biological Process) analysis against downregulated genes between 2wk-old and 80wk-old *C57BL/6J* mice; (C) GSEA analysis of downregulated pathway in response to the iron homeostasis; (D) Heatmap of cellular iron homeostasis related gene expression in TA muscle across five different aged (2wk-, 8wk-, 30wk-, 60wk- and 80wk-old); (E) qPCR analysis of *Tfr1* expression in four types of skeletal muscles (TA, EDL, Sol and Gas) between 2wk-old and 8wk-old *C57BL/6J* mice (n=5/group); (F) Representative western blot image of four types of skeletal muscles (TA, EDL, Sol and Gas) between 2wk-old and 8wk-old *C57BL/6J* mice (n=5/group); (G) Representative images of myofibers isolated from 2wk-old and 8wk-old *C57BL/6J* mice (n>50 myofibers from 5 mice/group). Immunofluorescence of Pax7 (red), Tfr1 (green), Ki67 (pink) and DAPI (blue) staining revealed Tfr1 is highly expressed in SCs at proliferative state (Ki67^+^) for 2wk-old mice but not 8wk-old adult mice; (H) Number of Ki67^+^ and Ki67^-^ SCs with different Tfr1 expression level (High, Inter and Low) per myofiber; (I) Number of Pax7^+^ SCs per myofiber; (J) Number of Ki67^+^ and Ki67^-^ SCs per myofiber. N.S.: not significant, \*\**p*<0.01, \*\*\**p* < 0.005, by 2-sided Student’s t-test. Data represent the mean ± SEM

### *Tfr1* is highly expressed in proliferative satellite cells

SC contributes to not only the skeletal muscle development but also postnatal myofiber formation and skeletal muscle regeneration*(12)*. However, the biological properties of the Tfr1-iron axis in SCs remain unexplored. Here, by performing single myofiber isolation and immunostaining, we observed that Tfr1 protein expression level was higher in SCs of 2wk-old mice than in SCs of 8wk-old mice (Figure 1H). Pax7, Ki67 and Tfr1 immunostaining of single myofibers was performed for both 2wk- and 8wk-old *C57BL/6J* mice. Approximately 70% of SCs on the myofibers of 2wk-old mice were Ki67^+^, compared with 8wk-old mice, and only ~7% of SCs turned into Ki67^+^. The Tfr1 expression level in SCs was highly correlated with the level of Ki67 expression in SCs. We quantified Tfr1 protein expression into three levels, High, Intermediate (Inter), and Low. Here, both Tfr1^High^/Ki67^+^ and Tfr1^Inter^/Ki67^+^ SCs were the dominant populations in young mice, while Tfr1^Low^/ Ki67^-^ SCs were barely detected. In contrast, it was completely the opposite in adult mice, whereas Tfr1^Low^/ Ki67^+^ SCs were the dominant population. In adult mice, both the Tfr1^High^/ Ki67^+^ and Tfr1^Inter^/ Ki67^+^ populations remained at low amounts (Figure 1I-J). The Tfr1 protein level was also determined in the SCs of aged mice (>80-wk old). Compared to SCs of adult mice, Tfr1 expression was almost undetectable in SCs of aged mice (Figure S3A). These results suggest that the Tfr1 protein level in SCs positively correlates with the cell proliferation status but negatively correlates with skeletal muscle ageing.

### *Tfr1*-mediated iron absorption is indispensable to SC proliferation and differentiation

Tfr1 protein is highly expressed in proliferative cells, while its expression is barely detected in quiescent SCs. To further prove this hypothesis, single myofibers were isolated and cultured *ex vivo* to induce SC proliferation and differentiation. In adult quiescent SCs (Pax7^+^/MyoD^-^), Tfr1 protein was expressed at low level. However, upon *ex vivo* culture, the Tfr1 protein level was dramatically induced in activated SCs (Pax7^+^/MyoD^+^, 24, 48 and 72-hr post-culture) (Figure S3B-C). This observation could be due to metabolic alterations during activation and proliferation, which require iron to support mitochondrial energy and glucose metabolism in newly activated SCs. To support the biological effect of iron on SC proliferation and differentiation, single myofibers were cultured with the DFO treatment to reduce extracellular iron availability to SCs. Upon 72-hr of culture, it resulted in a small cluster size, suggesting that iron is an essential component for SC proliferation (Figure S4A-B). DFO-mediated iron chelation also led to poor differentiation and myotube formation (Figure S4C-D). Furthermore, gradually increasing intracellular iron also significantly inhibited SC proliferation, cluster formation, differentiation and fusion to form mature myotubes (Figure S4E-H). All of these data indicate that Tfr1-mediated iron homeostasis is critical to support myoblast proliferation and differentiation.

### Tfr1 is required for maintaining SC homeostasis

To further understand the biological function of Tfr1 in SCs, SC-specific *Tfr1* knockout mice were generated by crossing mice carrying *Tfr1^fl/fl^* with *Pax7-CreER* transgenic mice. The genotype of experimental mice was *Pax7-CreER;Tfr1^fl/fl^*. *Tfr1-KO* mice and control littermates were denominated as *Tfr1^SC/KO^* and *Tfr1^SC/WT^*, respectively. The deletion was induced by intraperitoneal (*i.p*.)-injection of tamoxifen (TMX) for 7 consecutive days as described previously *(24)*. Single myofiber isolation from EDL of *Tfr1^SC/WT^* and *Tfr1^SC/KO^* was performed, followed by immunostaining of Pax7, MyoD, and Tfr1. Seven days post-injection (dpi) of tamoxifen (TMX), the number of Pax7^+^ SCs decreased in *Tfr1^SC/KO^* compared to *Tfr1^SC/WT^* (**Figure 2A-C**). Meanwhile, SCs with ablation of *Tfr1* did not express Ki67 or MyoD, indicating that *Tfr1*-deletion did not activate SCs (Figure 2A-C). Furthermore, the number of SC on a single myofiber was counted post-TMX injection (1, 4, 7, 10, 14, 21, and 30-dpi). Short term of TMX injection (1- and 4-dpi) did not lead to a change in the number of SCs, while the SC number gradually decreased since 7 dpi of TMX (Figure 2D). Deletion of *Tfr1* in SCs blocks TBI absorption, which may cause defects in SC proliferation and differentiation. To test this hypothesis, single myofibers from either *Tfr1^SC/WT^* or *Tfr1^SC/KO^* mice were cultured in horse serum-coated plates (flowing culture) or collagen-coated plates (attached culture) in the presence of 4-OH tamoxifen. Tamoxifen-induced deletion of *Tfr1* inhibited SC activation, proliferation and differentiation, as demonstrated by significantly decreased SC clusters, a reduced number of SCs in each cluster, failed myotube formation and a lower fusion index (Figure 2E-H). *Tfr1*-deletion inhibiting myoblast proliferation and differentiation was further tested in myoblasts bearing floxed *Tfr1*, whereas its deletion was induced by adenovirus expressing Cre recombinase. Both control and Cre-expressing adenovirus treated myoblasts were incubated with EdU-containing medium (10 μM) for 24-hr. The incorporation of EdU was significantly lower in the *Tfr1* deletion group (20%) than in the control group (60%) (Figure S5A-D)

**Figure 2.**
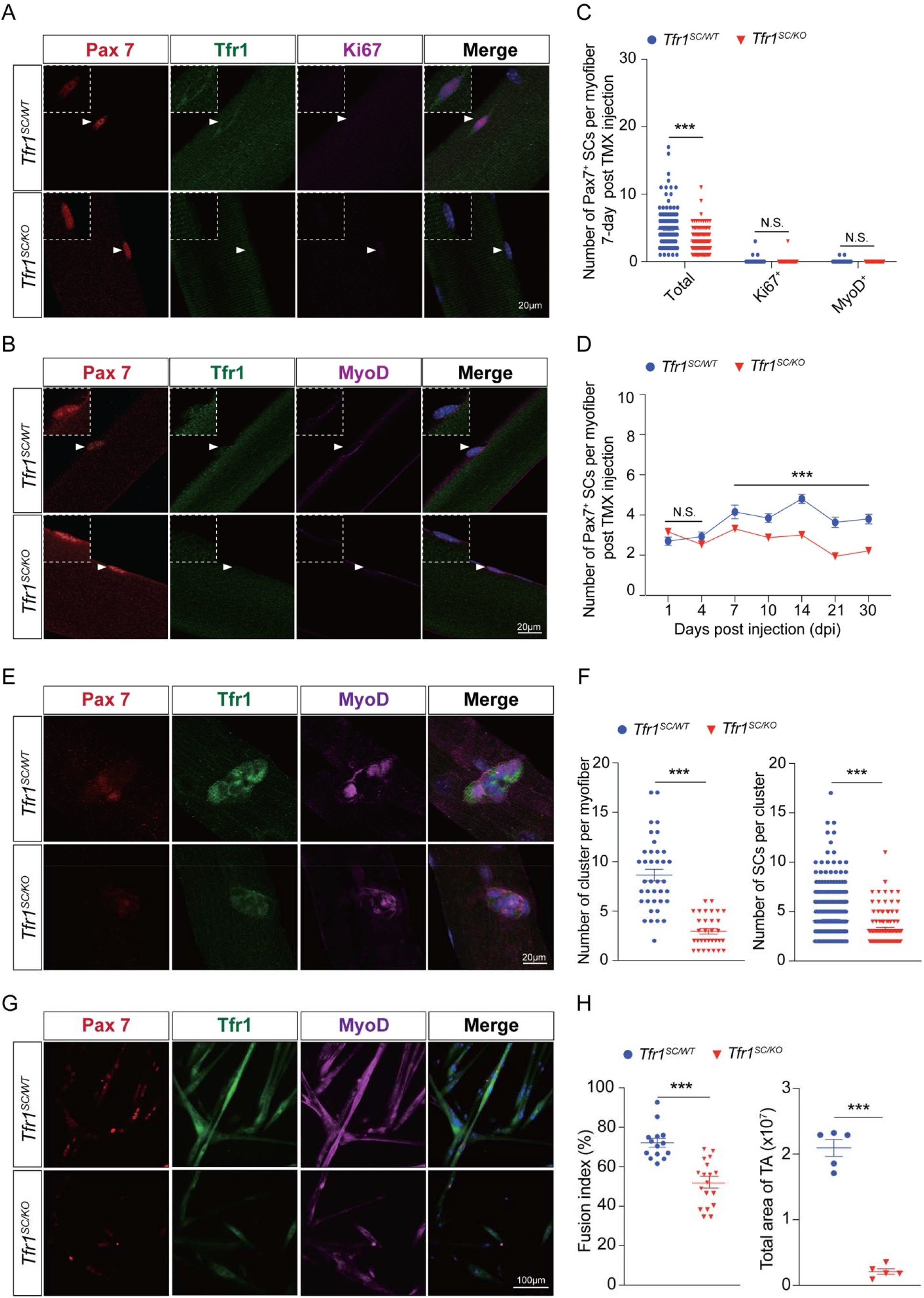
Genetic deletion of *Tfr1* in quiescent SCs abolishes the activation, proliferation and differentiation. (A) Representative images of myofibers isolated from *Tfr1^SC/WT^* and *Tfr1^SC/KO^* mice (n>50 myofibers from 5 mice/group). Immunofluorescence of Pax7 (red), Tfr1 (green), Ki67 (pink) and DAPI (blue) staining; (B) Representative images of myofibers isolated from *Tfr1^SC/WT^* and *Tfr1^SC/KO^* mice (n>50 myofibers from 5 mice/group). Immunofluorescence of Pax7 (red), Tfr1 (green), MyoD (pink) and DAPI (blue) staining; (C) Number of total, Ki67^+^ and MyoD^+^ SCs per myofiber between *Tfr1^SC/WT^* and *Tfr1^SC/KO^* mice; (D) Both *Tfr1^SC/WT^* and *Tfr1^SC/KO^* mice were administrated with tamoxifen on the same day. Number of Pax7^+^ SCs per myofiber was calculated at 1-, 4-, 7-, 10-, 14-, 21- and 30-day post tamoxifen-induced *Tfr1* deletion (n = 5 mice/group/time point); (E) Representative images of SC clusters on myofiber from *Tfr1^SC/WT^* and *Tfr1^SC/KO^* mice *ex vivo* cultured for 72 hrs (n>50 myofiber). Immunofluorescence of Pax7 (red), Tfr1 (green), MyoD (pink) and DAPI (blue) staining (n >20 myofibers from 7 mice/group); (F) Number of SC clusters per myofiber and number of Pax7^+^ SC per cluster (n>50 myofibers from 5 mice/group); (G) Representative images of differentiated myotubes from SCs on myofiber isolated from *Tfr1^SC/WT^* and *Tfr1^SC/KO^* mice (n>10 myofibers from 5 mice/group); (H) Summary of fusion index of SCs on myofiber differentiated in DMEM supplemented with 2% horse serum (n>10 myofibers from 5 mice/group). N.S.: not significant, ***P < 0.005, by 2-sided Student’s t-test. Data represent the mean ± SEM.

### *Tfr1*-deletion disrupts skeletal muscle regeneration *via* the inhibition of SC proliferation

The biological function of Tfr1 in skeletal muscle was investigated before showing that *Tfr1*-ablation is lethal if there is no additional ferric iron administration, accompanied by systemic metabolic disorders*(22)*. To understand the biological function of Tfr1 in the exercise system regarding skeletal muscle development, growth and regeneration, we utilized mice with conditional deletion of *Tfr1* in SCs. 14 dpi of TMF, TA muscle was injured by intramuscular injection of cardiotoxin (CTX) and harvested for further analysis upon the completion of regeneration (Figure 3A). We observed poor regeneration with Tfr1 dysfunction in SCs, showing a clear muscular atrophy but clearly no change in body weight (Figure 3B-C). To assess the biological function of Tfr1 during TA muscle regeneration, the number of Pax7^+^ SCs and eMyHC^+^ myotubes (newly formed myotubes) was counted. Upon CTX-induced injury (5 and 9 dpi), Pax7^+^ SCs were highly proliferated in *Tfr1^SC/WT^* mice, while they were barely detected in *Tfr1^SC/KO^* mice on TA sections (Figure 3D-E). Low SC numbers were further confirmed by *Pax7* mRNA expression indicating that *Tfr1* deletion led to the depletion of Pax7^+^ SCs upon skeletal muscle injury (Figure S6A). *Tfr1* KO in SCs also decreased eMyHC^+^ myotube formation at 5 dpi (Figure 3F) while there was a dramatic increase in eMyHC^+^ myotubes at 9 dpi (Figure 3G), which may be due to a robust induction of myogenic transcription factor, *MyoD* expression (Figure S6A). Upon the completion of regeneration, *Tfr1* KO caused almost complete depletion of SCs in the TA, followed by atrophy and fibrosis (Figure 3H-K and Figure S6C-E).

**Figure 3.**
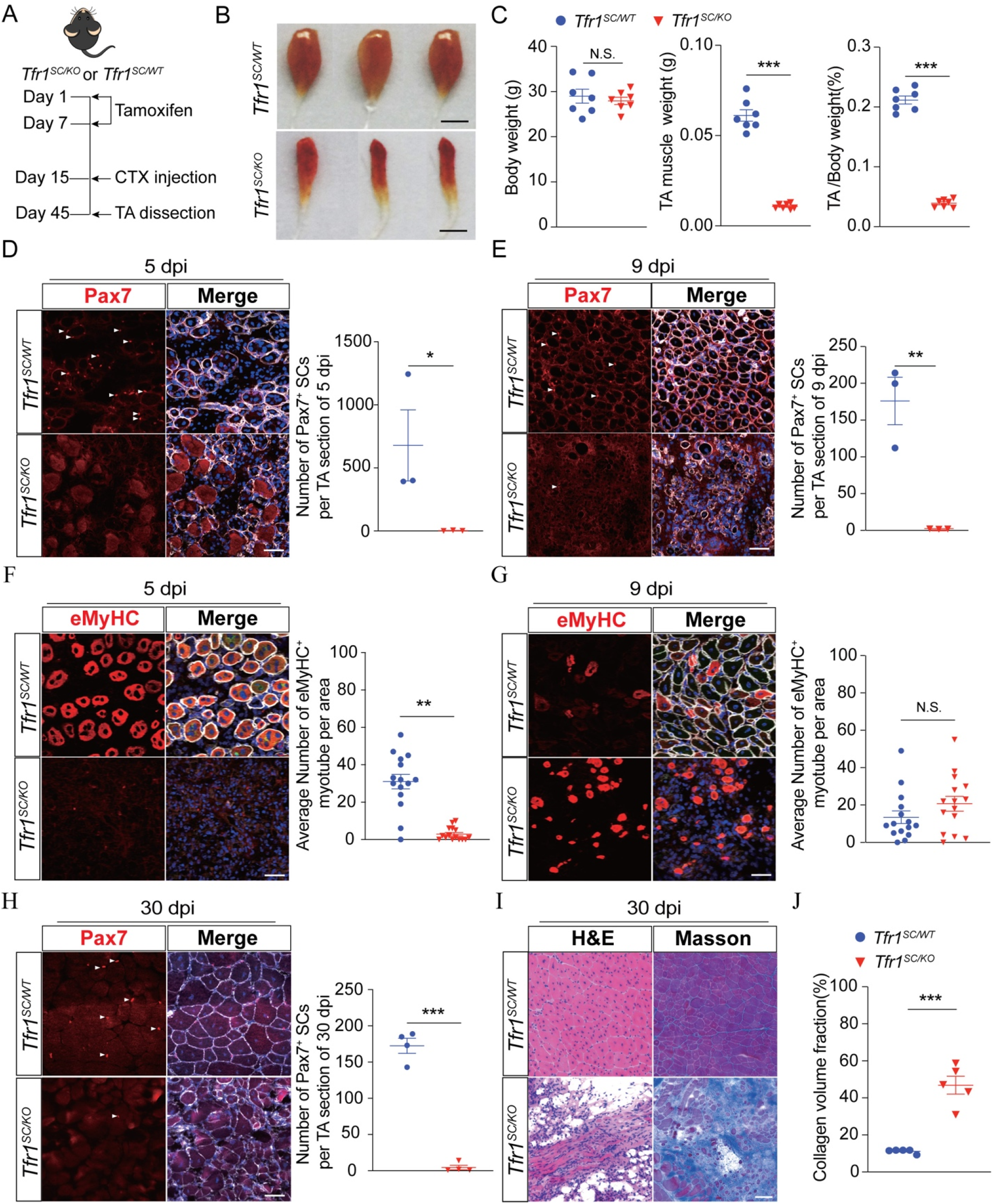
*Tfr1*-ablation in SCs blocks skeletal muscle regeneration. (A) Timeline characterizing skeletal muscle regeneration upon tamoxifen-induced *Tfr1*-ablation in SCs; (B) *SC/WT SC/KO* Representative image of TA upon completion of regeneration between *Tfr1^SC/WT^* and *Tfr1^SC/KO^* mice at 30 dpi; (C) Summary of body weight, TA weight and ratio of TA and body weight between *Tfr1^SC/WT^* and *Tfr1^SC/KO^* mice upon completion of regeneration at 30 dpi (n=7/group); (D) Representative images of TA section from *Tfr1^SC/WT^* and *Tfr1^SC/KO^* mice (n=3 mice/group). Immunofluorescence of Pax7 revealed a decrease in the number of Pax7^+^ SCs (arrowheads) and number of Pax7^+^ SCs per TA muscle section at 5 dpi (right of immunostaining images); (E) Representative images of TA section from *Tfr1^SC/WT^* and *Tfr1^SC/KO^* mice (n=3 mice/group). Immunofluorescence of Pax7 revealed a decrease in the number of Pax7^+^ SCs (arrowheads) and number of Pax7^+^ SCs per TA muscle section at 9 dpi (right of immunostaining images); (F) Immunofluorescence of eMyHC^+^ myotubes after CTX injury (5 dpi) and number of eMyHC^+^ myotubes per TA section a rea at 5 dpi (right of immunostaining images); (G) Immunofluorescence of eMyHC^+^ myotubes after CTX injury (5 dpi) and number of eMyHC^+^ myotubes per TA section area at 9 dpi (right of immunostaining images); (H) Immunofluorescence of Pax7 revealed a decrease in the number of Pax7^+^ SCs (arrowheads) and number of Pax7^+^ SCs per TA muscle section at 30 dpi (right of immunostaining images, n=4 mice/group); (I) Representative images of TA muscles from *Tfr1^SC/WT^* and *Tfr1^SC/KO^* mice with H.E. and Masson staining upon completion of CTX-induced regeneration (30 dpi, n=6 mice/group); (J) Summary of collagen volume fraction between *Tfr1^SC/WT^* and *Tfr1^SC/KO^* mice completion of CTX-induced regeneration at 30 dpi. N.S.: not significant, *P < 0.05, **P < 0.01, ***P < 0.005, by 2-sided Student’s t-test. Data represent the mean ± SEM.

### SC Tfr1 is essential to maintain the skeletal muscle microenvironment and regeneration

To precisely understand how SC-specific knockout of Tfr1 affects the skeletal muscle microenvironment, RNAseq was performed to assess the gene expression profile in the TA muscle between *Tfr1^SC/WT^* and *Tfr1^SC/KO^* mice before or after injury. Approximately 8478 differentially expressed genes were identified among four groups (Figure S7A). Gene clustering and principal coordinates analysis (PCoA) showed that *Tfr1^SC/KO^* mice with CTX injection exhibited a distinct molecular signature from the other three groups, which had similar molecular *SC/WT* signatures (Figure 4B). Thus, we focused on DEGs and functional enrichment between *Tfr1^SC/WT^* and *Tfr1^SC/KO^* mice post-CTX-induced injury. Among DEGs, 3596 genes were upregulated, while 4882 genes were downregulated (Figure 4C). Gene ontology of biological process gene set analysis identified genes that were majorly involved in dysregulation of immune balancing and metabolic homeostasis. This was represented by upregulated genes that were associated with macrophage activation, macrophage-derived foam cell differentiation, lipid biosynthetic process and collagen biosynthetic process, while downregulated genes were involved in mitochondrial respiration chain complex assembly, TCA cycle, muscle cell differentiation and fatty acid beta-oxidation (Figure 4D and Figure S7B-E). The enriched biological functions were further confirmed by GSEA (Figure 4E and 4H). Here, we found that in regenerated TA muscle, *Tfr1* knockout resulted in macrophage infiltration, which was assessed by flow cytometry by detecting M1 (0.17% vs 0.012%) and M2 (0.88% vs 872E-3%) macrophages by their membrane markers, e.g. Cd86, Cd206 and Cd163 (Figure 4F). *Cd86, Cd206* and *Cd163* mRNA expression was robustly increased only in regenerated TA of *Tfr1^SC/KO^* mice but not in other groups (Figure 4G). Defective muscle regeneration was also accompanied by extracellular collagen biosynthesis accumulation, with the upregulation of collagen biosynthesis and accumulation related genes (Figure 4I-L and Figure S7C). This would further interrupt exercise activity by reducing the running time and distance (Figure 4K-L).

**Figure 4.**
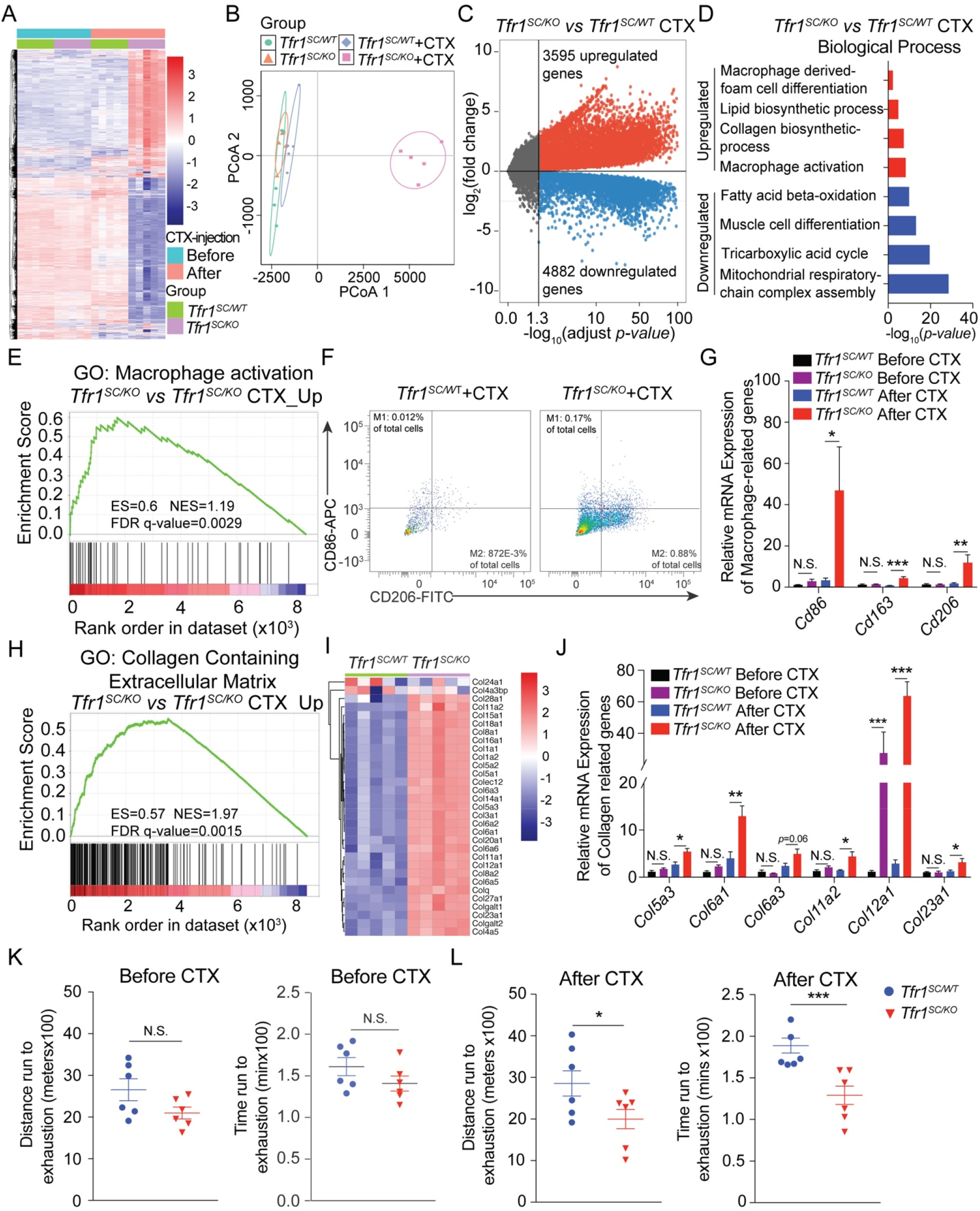
SCs-*Tfr1* deletion in TA of *Tfr1^SC/KO^* mice results in skeletal muscle dysfunction. (A) Heatmap of mRNA expression profile in TA muscle from adult *Tfr1^SC/WT^* and *Tfr1^SC/KO^* mice before or after CTX-induced regeneration at 30 dpi (n=5/group); (B) PCoA of transcriptome from TA muscle in adult *Tfr1^SC/WT^* and *Tfr1^SC/KO^* mice before or after CTX-induced regeneration (n=5/group); (C) Volcano plot of differentially expressed genes in TA muscle from adult *Tfr1^SC/WT^* and *Tfr1^SC/KO^* mice after CTX-induced regeneration; (D) GO (Biological Process) analysis of DEGs for both upregulated and downregulated genes; (E) GSEA analysis of macrophage activation pathway between *Tfr1^SC/WT^* and *Tfr1^SC/KO^* mice upon CTX-induced regeneration at 30 dpi; (F) Flow cytometry analysis of the percentage of the CD206^+^/CD86^+^ macrophage in total cells obtained from TA of *Tfr1^SC/WT^* and *Tfr1^SC/KO^* mice after CTX-induced injury at 30 dpi; (G) qPCR analysis of *Cd86, Cd163* and *Cd206* mRNA expression in TA muscle from adult *Tfr1^SC/WT^* and *Tfr1^SC/KO^* mice before or after CTX-induced injury (n=5/group); (H) GSEA analysis of collagen containing extracellular matrix pathway between *Tfr1^SC/WT^* and *Tfr1^SC/KO^* mice upon CTX-induced regeneration at 30 dpi; (I) Heatmap for collagen matrix related gene expression profile between *Tfr1^SC/WT^* and *Tfr1^SC/KO^* mice after CTX-induced injury at 30 dpi (n=5); (J) qPCR analysis of *Col5a3, Col6a1, Col6a3, Col11a2, Col12a1*, and *Col23a1* mRNA expression in TA muscle from adult *Tfr1^SC/WT^* and *Tfr1^SC/KO^* mice before or after CTX-induced injury at 30 dpi (n=5/group); (K-L) Tread mill running distance and running time to exhaustion for *Tfr1^SC/WT^* and *Tfr1^SC/KO^* mice before and after regeneration at 30 dpi. N.S.: not significant, *P < 0.05, **P < 0.01, ***P < 0.005, by 2-sided Student’s t-test. Data represent the mean ± SEM.

### *Tfr1*-deletion in SCs leads to the dysregulation of lipid and iron metabolism in skeletal muscle

SC-specific deletion of *Tfr1* results in dysregulation of local lipid and energy metabolism (Figure 4D). GSEA analysis further confirmed that adipogenesis indeed occurred in the TA of *Tfr1^SC/KO^* mice (Figure 5A). Adipogenesis-related genes such as *Fasn* and *Adipoq* were significantly induced. However, the expression of the fatty acid uptake gene, *Cd36* expression was not changed, indicating that a local *de novo* lipogenesis instead of external fatty acid uptake contributed to the lipid accumulation in TA muscle of *Tfr1^SC/KO^* mice (Figure 5B). Lipid accumulation in TA muscle was visualized by Oil Red O (ORO) staining and immunofluorescent staining (IF) of Perilipin and Laminin B2. On TA cryosection, a large amount of lipid droplets as well as Perilipin^+^ areas were observed only in TA of *Tfr1^SC/KO^* mice but not in *Tfr1^SC/WT^* mice at 30 dpi (Figure 5C). Other than dysregulation of genes associated with lipid metabolism, most mitochondrial thermogenesis and iron metabolism-related genes were dramatically downregulated, such as *Pgc1α, Cox7a1* and *Cox8b* for mitochondrial thermogenesis and *Tfr1, Slc11a2, Slc40a1* and *Fth1* for iron metabolism, except for a moderate upregulation of *Ftl* (Figure 5D). As reported before, Tfr1 is indispensable for iron assimilation by skeletal muscle*(25)*. Muscular dysfunction of the Tfr1 gene results in systemic metabolic disorders, such as iron deficiency in muscle and liver, as well as systemic glucose and lipid disorders*(25)*. In contrast, atrophied skeletal muscle of *Tfr1^SC/KO^* mice was not iron deficient instead of having a large amount of labile iron accumulation (Figure 5E). Furthermore, consistent with mitochondrial gene expression, transmission electron microscopy revealed swollen mitochondria with irregular or absent cristae (Figure 5E).

**Figure 5.**
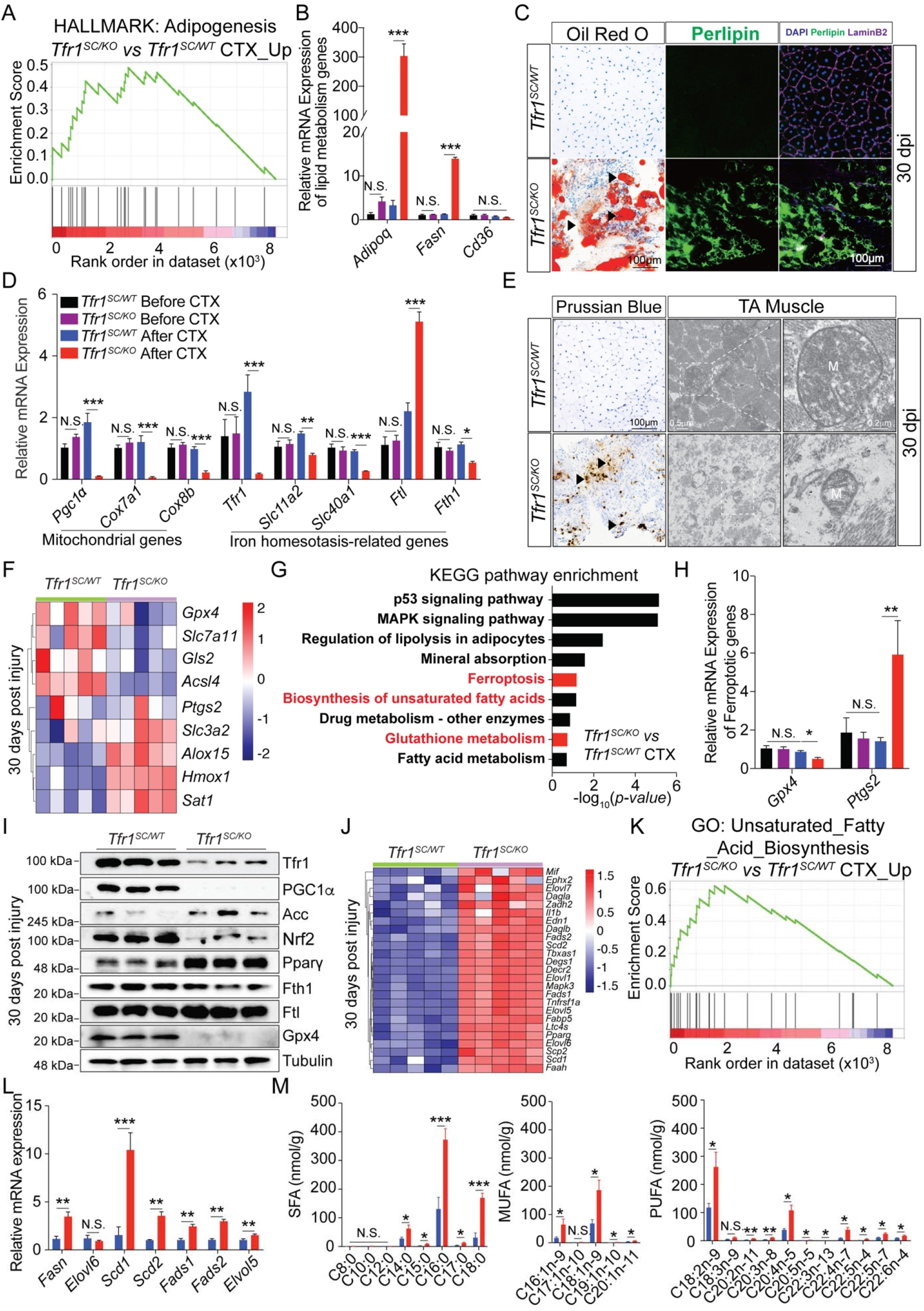
Dysregulation of lipid and iron metabolism activates ferroptosis in injured TA of *Tfr1^SC/KO^* mice. (A) GSEA analysis of adipogenesis pathway between *Tfr1^SC/WT^* and *Tfr1^SC/KO^* mice upon CTX-induced regeneration at 30 dpi; (B) qPCR analysis of *Adipoq, Fasn* and *Cd36* mRNA expression in TA muscle from adult *Tfr1^SC/WT^* and *Tfr1^SC/KO^* mice before or after CTX-*SC/WT* induced injury (n=5/group); (C) Representative images of TA sections from *Tfr1^SC/WT^* and *Tfr1^SC/KO^* mice upon CTX-induced injury at 30 dpi (n=5/group). Oil Red O (ORO) staining and perilipin (green) and Laminin B2 (pink) immunofluorescent staining revealed adipogenesis and lipid accumulation in TA of *Tfr1^SC/KO^* mic; (D) qPCR analysis of *Pgc1α, Cox7a1*, and *Cox8b* (mitochondrial genes), *Tfr1, Slc11a2, Slc40a1, Ftl* and *Fth1* (iron homeostasis related genes) mRNA expression in TA muscle from adult *Tfr1^SC/WT^* and *Tfr1^SC/KO^* mice before or after CTX-induced injury (n=5/group); (E) Representative images of TA section with Prussian Blue staining *SC/WT* (n=5/group) and transmission electron microscope images for TA section from adult *Tfr1^SC/WT^* and *Tfr1^SC/KO^* mice after CTX-induced injury at 30 dpi; (F) Heatmap of ferroptosis-related gene expression in TA from adult *Tfr1^SC/WT^* and *Tfr1^SC/KO^* mice after CTX-induced injury at 30 dpi (n=5/group); (G) KEGG pathway enrichment analysis of upregulated genes in TA from adult *Tfr1^SC/WT^* and *Tfr1^SC/KO^* mice after CTX-induced injury at 30 dpi; (H) qPCR analysis of *Gpx4* and *Ptgs2* expression in TA of adult *Tfr1^SC/WT^* and *Tfr1^SC/KO^* mice before or after CTX-induced injury (n=5/group); (I) Representative western blot images of TA muscle between *Tfr1^SC/WT^* and *Tfr1^SC/KO^* mice after CTX-induced injury at 30 dpi; (J) Heatmap of unsaturated fatty acid biosynthesis-related gene expression in TA from adult *Tfr1^SC/WT^* and *Tfr1^SC/KO^* mice after CTX-induced injury at 30 dpi (n=5/group); (K) GSEA analysis of unsaturated fatty acid biosynthesis pathway between *Tfr1^SC/WT^* and *Tfr1^SC/KO^* mice upon CTX-induced regeneration at 30 dpi; (L) qPCR analysis of *Fasn, Elvol5, Elvol6, Scd1, Fads1* and, *Fads2* expression in TA of adult *Tfr1^SC/WT^* and *Tfr1^SC/KO^* mice after CTX-induced injury at 30 dpi (n=5/group); (M) SFU, MUFA and PUFA level (nmol/g) in TA of adult *Tfr1^SC/WT^* and *Tfr1^SC/KO^* mice after CTX-induced injury at 30 dpi (n=4/group). N.S.: not significant, *P < 0.05, **P < 0.01, ***P < 0.005, by 2-sided Student’s t-test. Data represent the mean ± SEM.

### *Tfr1*-deletion activates ferroptosis in skeletal muscle upon muscular regeneration

Regenerated skeletal muscle of *Tfr1^SC/KO^* mice had upregulated adipogenesis and iron accumulation in TA muscle. To profile new gene candidates between *Tfr1^SC/WT^* and *Tfr1^SC/KO^* mice at 30 dpi, we discovered a group of ferroptopic genes were dysregulated (e.g. *Gpx4, Slc7a11, Acsl4*, and *Hmox1*), along with the KEGG pathway enrichment, including ferroptosis, biosynthesis of unsaturated fatty acid and glutathione metabolism (Figure 5F-G). Consistent with previous observations in ferroptosis, *Gpx4*, a glutathione-dependent peroxidase was downregulated and *Ptgs2* (cyclooxygenase-2: Cox-2), an enzyme converting arachidonic acid (AA) to prostaglandin endoperoxide H2 was upregulated (Figure 5H). The activation of ferroptosis in skeletal muscle was further confirmed by measuring selective biomarker protein levels. This was shown by the induced protein levels of Acc and Pparγ proteins and the decreased protein levels of Tfr1, PGC1α, Nrf2, Gpx4 and Fth1, leading to the observation of adipogenesis and dysregulated iron metabolism, respectively, contributing to the activation of ferroptosis. Further analysis demonstrated that fatty acid biosynthesis was accompanied by the unsaturated fatty acid biosynthesis (Figure 5J-K). The critical enzymes in the unsaturated fatty acid biosynthesis pathway, e.g., *Fasn, Elvol5, Scd1, Scd2, Fads1*, and *Fads2* were upregulated, followed by accumulation of saturated and unsaturated fatty acids (Figure 5L-M, and Figure S8A-C).

### Ferroptosis decreases muscular regeneration

To map the time-point of ferroptosis occurrence, injured TA muscle from *Tfr1^SC/WT^* and *Tfr1^SC/KO^* mice at different time points (5, 9, and 15 dpi) was assessed (Figure 6A). The TA/BW, *SC/WT* as well as *Gpx4, Slc3a2* and *Ptgs2* expression, was not significantly different between *Tfr1^SC/WT^* and *Tfr1^SC/KO^* mice at 5 dpi (Figure 6B and Figure S9A-G). However, the expression of iron homeostasis related genes such as *Slc11a2, Slc39a14, Fth1* and *Ftl*, was highly upregulated at 5 dpi in *Tfr1^SC/KO^* mice, suggesting that increased iron absorption may occur as early as at 5 dpi (Figure S9A-G). Starting at 9 dpi, TA/BW started to decrease, followed by downregulation of *Gpx4* and upregulation of *Ptgs2, Slc39a14*, and *Hmox1* expression, so as to the TA at 15 dpi, except for decreased *Fth1* expression that is associated with oxidization of ferrous iron and iron storage in ferritin (Figure 6B and Figure S9A-G). Iron accumulation and lipid droplets could be observed in the TA muscle of *Tfr1^SC/KO^* mice starting at 9 dpi (Figure 6C). During muscle regeneration, labile iron in regenerative TA muscle was derived from increased non-heme iron absorption possibly *via* Slc39a14 and iron recycling from myoglobin from dead muscle cells but failed to be utilized upon muscular regeneration.

**Figure 6.**
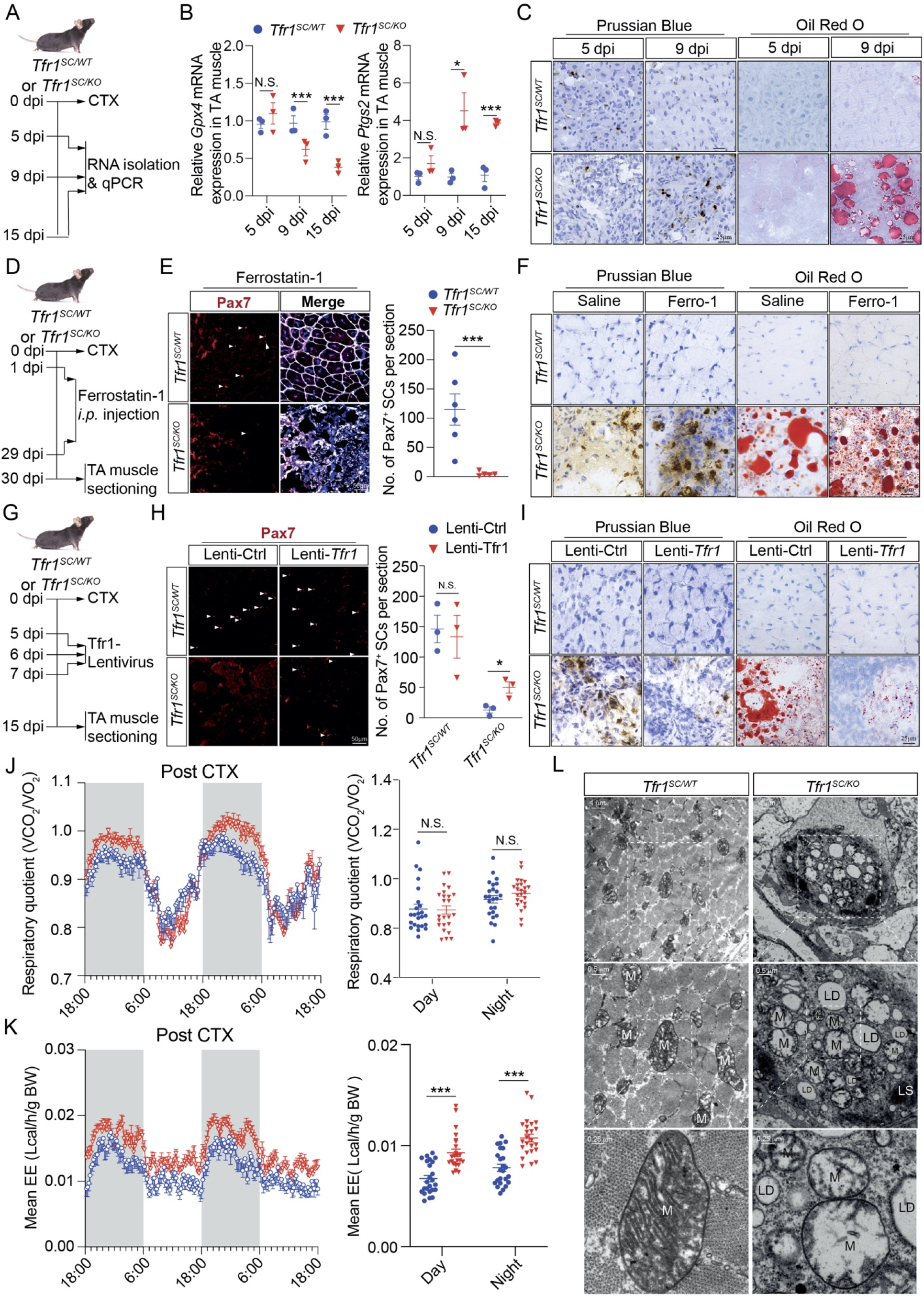
Ferroptosis in TA of *Tfr1^SC/KO^* mice prevents skeletal muscle regeneration. (A) Timeline characterizing the activation of ferroptosis in TA muscle of *Tfr1^SC/WT^* and *Tfr1^SC/KO^* mice after CTX-induced injury; (B) qPCR analysis of *Gpx4* and *Ptgs2* expression in TA from adult *Tfr1^SC/WT^* and *Tfr1^SC/KO^* mice after CTX-induced injury at 5, 9 and 15 dpi; (C) Representative images of TA section with Prussian Blue and ORO staining from *Tfr1^SC/WT^* and *Tfr1^SC/KO^* mice after CTX-induced injury at 5 and 9 dpi; (D) Timeline characterizing the effect of Ferrostatin-1 to inhibit ferroptosis in TA muscle from *Tfr1^SC/WT^* and *Tfr1^SC/KO^* mice; (E) Representative images of TA section immunostaining for Pax7 (red) and Laminin B2 (white). Number of Pax7^+^ SCs per section (right); (F) Representative images of TA section with Prussian Blue and ORO staining from adult *Tfr1^SC/WT^* and *Tfr1^SC/KO^* mice after CTX-induced injury between saline and Ferrostatin-1 *i.p*. injection at 30 dpi; (G) Timeline characterizing the effect of lenti-Tfr1 to inhibit ferroptosis in TA muscle from *Tfr1^SC/WT^* and *Tfr1^SC/KO^* mice; (H) Representative images of TA section immunostaining for Pax7 (red) and Laminin B2 (white) between lenti-Ctrl and lenti-Tfr1 intramuscular injection at 30 dpi. Number of Pax7^+^ SCs per section (right); (I) Representative images of TA section with Prussian Blue and ORO staining from adult *Tfr1^SC/WT^* and *Tfr1^SC/KO^* mice after CTX-induced injury between lenti-Ctrl and lenti-Tfr1 intramuscular injection at 15 dpi; (J) Respiratory exchange rate (VCO_2_/VO_2_) and energy expenditure were monitored over a 48-h period for *Tfr1^SC/WT^* and *Tfr1^SC/KO^* mice after CTX-induced injury at 15 dpi (n = 8 mice/group); (L) Representative transmission electron microscope image of TA muscle samples from adult *Tfr1^SC/WT^* and *Tfr1^SC/KO^* mice after CTX-induced injury at 30 dpi. N.S.: not significant, *P < 0.05, ***P < 0.005, by 2-sided Student’s t-test. Data represent the mean ± SEM.

Next, we asked whether administration of a ferroptosis inhibitor would reverse the ferroptosis induced muscular hypotrophy. Ferrostatin-1 (Ferro-1), a ferroptosis inhibitor to eliminate lipid peroxidation, was *i.p*.-injected upon intramuscular administration of CTX to TA muscle for 30 consecutive days (Figure 6D). However, Ferro-1 did not rescue ferroptosis-induced SC/muscle cell death or reduce labile iron accumulation and lipid droplet formation (Figure 6E-F). Instead, administration of lentivirus-expressing mouse Tfr1 protein could partially reverse ferroptosis-induced SCs/muscle cell death by decreasing the labile iron accumulation and lipid biosynthesis (Figure 6G-I).

### Metabolic adaptation of ferroptosis via mitochondrial stress in skeletal muscle

Skeletal muscle-specific deletion of *Tfr1* leads to growth retardation and systemic metabolic disorder (lipid and amino acid) in both muscle and liver*(18)*. However, for our model, data from mice kept in metabolic cages presented a significantly higher energy expenditure (EE) for *Tfr1^SC/KO^* mice than that of *Tfr1^SC/WT^* mice but no difference in the ratio of O_2_ consumption and CO_2_ production, meaning that an adaptive alteration of systemic metabolism, especially the induction of EE, was probably not due to the change in substrate preference and/or whole-body fuel metabolism (Figure 6J-K). Through the transmission electron microscopy, we observed a lysosomal structure containing dead mitochondria (M) without any cristae structure and lipid droplets (LD) in *Tfr1^SC/KO^* mice (Figure 6L), with higher levels of Fgf21 but lower levels of Trp53 and mitochondrial complex protein (Complexes I, II, III, and V) in the TA of *Tfr1^SC/KO^* mice (Figure S9). Increased Fgf21 protein may be due to mitochondrial stress*(26, 27)*. To eliminate potential endocrinological regulation of systemic metabolism and thermogenesis by Fgf21, iron metabolism and thermogenesis-related genes were determined in liver, iBAT, iWAT *SC/WT SC/KO* and eWAT, showing that no difference in their gene expression between *Tfr1^SC/WT^* and *Tfr1^SC/KO^* mice was detected (Figure S10A-H, Supporting Information). However, Glut4 protein levels were significantly induced in both iWAT and eWAT but not in iBAT (Figure S10F and S10H, Supporting Information). These results demonstrated that increased EE was not due to the metabolic alteration in distal tissues instead of the mitochondrial stress with the occurrence of ferroptosis in skeletal muscle.

### Skeletal muscle ageing accompanied by a Tfr1 and Slc39a14 functional switch to mediate labile iron accumulation and the activation of ferroptosis

Skeletal muscle ageing is known as sarcopenia with the loss of muscle mass and function, which may be due to multifactorial conditions, e.g. imbalance between protein synthesis and degradation*(28, 29)*, reduced number of satellite cells*(30)* and increased production of ROS*(31, 32)*. It has been reported that aged skeletal muscle has more labile iron*(33, 34)*. To analyze 2445 DEGs between aged and young mice and 1333 DEGs between *Tfr1^SC/WT^* and *Tfr1^SC/KO^* mice, 2203 common biomarkers were identified from two datasets. Among them, 72 genes were universally upregulated, and 132 genes were downregulated (Figure 7A-B). Through the KEGG pathway enrichment analysis for these common genes between the two groups, ferroptosis, glutathione metabolism and fatty acid biosynthesis were the top candidate pathways, suggesting that skeletal muscle ageing may be accompanied by ferroptosis. Ferrozine assay to assess the serum and TA muscle non-heme iron showed that compared to the young mice, aged mice had significantly higher iron levels (Figure 7D). Aged TA muscle expressed lower Tfr1, Gpx4 and Fth1 but higher Slc39a14, which mimics the gene expression pattern in *Tfr1^SC/KO^* mice (TA total protein, Figure 7E). Most importantly, TA membrane Tfr1 protein was decreased to undetected levels but with significantly higher expression levels of Scl39a14 (Figure 7E). These observations indicated that Slc39a14 facilitates NTBI absorption in aged skeletal muscle causing iron accumulation and ferroptosis. Meanwhile, Ferro-1 as a ferroptosis inhibitor, was *i.p*.-injected into aged mice upon intramuscular injection of CTX to induce injury and regeneration. Thirty days post-injection indeed revealed improved running capacity, such as running time and distance, but did not reach the statistical significance (Figure 7F).

**Figure 7.**
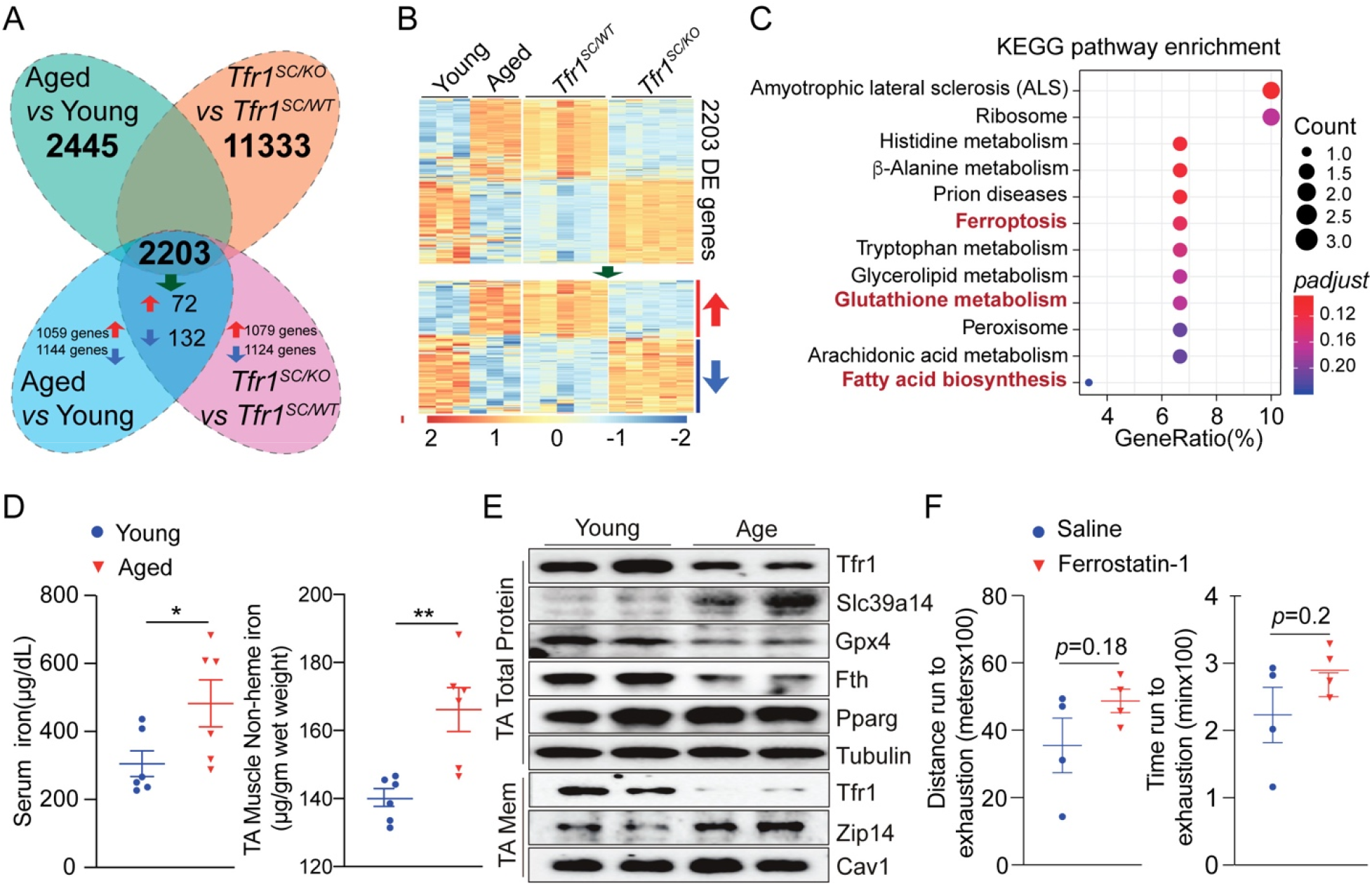
Slc39a14-mediated iron absorption and labile iron accumulation induces ferroptosis in aged skeletal muscle. (A) Venn diagraph showing the overlapping genes between Aged/Young and *Tfr1^SC/WT^*/*Tfr1^SC/KO^* samples; (B) Heatmap of overlapping gene expression profile for Aged/Young group (n=3) and *Tfr1^SC/WT^/Tfr1^SC/KO^* group (n=5); (C) KEGG pathway enrichment analysis of upregulated common genes identified ferroptosis-related genes highly expressed in TA muscle of aged mice; (D) Serum and total TA muscle non-heme iron from young (8wk-old) and aged (80wk-old) *C57BL/6J* mice; (E) Representative western blot image of total and membrane protein of TA muscle from young (8wk-old) and aged (80wk-old) *C57BL/6J* mice; (F) Treadmill running distance and running time to exhaustion for aged (80wk-old) *C57BL/6J* mice with CTX-induced injury followed by *i.p*. injection of either saline or Ferrostatin-1 for 30 days. .S.: not significant, *P < 0.05, **P < 0.01, by 2-sided Student’s t-test. Data represent the mean ± SEM.

## Discussion

Programmed cell death, such as apoptosis, autophagy, and necrosis, is stimulated by external factors in response to muscle injury. Dysregulation of these processes results in muscle loss and sarcopenia. Here, in our regeneration defect model, we are surprised to identify a new form of cell death, which has not been reported elsewhere in skeletal muscle. Newly defined iron-dependent cell death, also named ferroptosis is activated during muscle regeneration with *Tfr1*-deletion in SCs. This event is coupled with the upregulation of *Slc39a14* expression to uptake NTBI, unsaturated fatty acid biosynthesis, and decreased expression of anti-ferroptosis biomarkers such as Gpx4 and Nrf2. Most importantly, the scenario of Tfr1-Slc39a14-iron axis is recapitulated in aged skeletal muscle of rodents, which provides physiological relevance of the Tfr1-Scl39a14-iron axis and ferroptosis in skeletal muscle.

Iron homeostasis is indispensable to the proper function of skeletal muscle and postnatal regeneration, reflected from the importance of iron in mitochondrial respiration, ATP production, muscle contraction and exercise capacity. This is partially due to the activity of the mitochondrial electron transport chain and mitochondrial clearance*(35)*. These observations could be manipulated in murine models fed with iron-deprived diets, but muscular iron deficiency in patients is usually accompanied by secondary diseases, such as congestive heart failure and chronic obstructive pulmonary disease*(36, 37)*. In terms of the developmental essentiality of iron, numerous studies have demonstrated that iron deficiency during pregnancy or the early stage of development causes growth retardation. However, a high systemic iron level is detrimental to the host, especially at advanced age. Genome-wide association studies (GWAS) identified the association between healthy longevity and iron traits, e.g., serum iron, transferrin level, and transferrin saturation*(38)*. Furthermore, individuals at advanced age experience iron accumulation in multiple organs, such as the brain, skeletal muscle and liver*(39)*. In contrast to iron deficiency, iron overloading or accumulation in tissue leads to increased oxidative stress by producing highly toxic hydroxyl radicals through the Fenton reaction. High iron content, especially the non-heme iron (NHI), is associated with decreased muscle mass in both elderly human and aged rat skeletal muscle*(19, 40)*. Evidence from different studies are consistent including ours that aged skeletal muscle has a phenotype of elevated NHI content, impaired muscle function and muscular atrophy. In the *ex vivo* culture system, a high iron content decreases the single myofiber survival rate, reduces SC cluster formation and prohibits myoblast differentiation. However, the mechanisms causing iron accumulation and muscle loss remain unknown and need further exploration.

To delve into the pathogenic effect of iron and identify the potential mediator, we used multiple approaches and models to explore. As reported previously*(40)*, we also noted declined *Tfr1* mRNA and protein in both skeletal muscles and SCs of older mice but with higher NHI levels in skeletal muscle*(39)*. We also found that the membrane Tfr1 protein is almost undetectable in aged mice (>80-wk old) compared to young mice (~8-wk old). To profile the expression of other iron absorption related genes, *Slc11a2* and *Slc39a14* expression were upregulated in TA muscle of 80wk-old mice (Figure 1D). However, Slc11a2-mediated TBI iron absorption in peripheral tissue relies on the membrane Tfr1 protein. Our work for the first-time identifies the phenomenon of Tfr1/Slc39a14 expression switching in aged skeletal muscle. More specifically, Slc39a14 expression is induced and its protein is enriched on the cellular membrane to facilitate NTBI absorption to replace Tfr1. This corresponds to the NHI accumulation in aged TA muscle. Our work provided evidence to interpret how labile iron accumulated in the aged skeletal muscle.

Under physiological conditions, ferric iron in the form of transferrin-bound iron (TBI) recognized by membrane Tfr1 is absorbed in peripheral tissues, such as liver, adipose tissue, skeletal muscle, and bone marrow*(23, 41, 42)*. Multiple studies have shown that *Tfr1*-deficiency results in functional disorder and even death. Mice with *Tfr1* conditional knockout in hematopoietic stem cells died within one week after birth*(43)*. Our previous study demonstrated that Tfr1 regulates adipocyte thermogenesis and cell fate determination*(44)*. Adipocytes with *Tfr1* ablation exhibit reduced thermogenic capacity and beigeing potential*(44)*. Meanwhile, an alpha-skeletal actin (HSA)-driven Cre recombinase expression results in embryonic dysfunction of *Tfr1*, disrupting iron homeostasis and leading to the muscle growth retardation*(18)*. This may be associated with dysfunction of heme-containing myoglobin synthesis and energy metabolism in muscle and liver. However, postnatal function of Tfr1 in skeletal muscle could not be well-explored by utilizing this model as postnatal growth and regeneration of skeletal muscle rely on the activity of SCs. Thus, to solve the discrepancy, mice with conditional knockout of *Tfr1* in SCs were generated. In additional to regeneration defects, during muscle regeneration, we observed that failed SCs activation and muscle regeneration, myoglobin degradation by heme oxygenase 1 (Hmox1) and upregulation of *Slc39a14* expression may be three key factors contributing to labile iron accumulation, which may also be followed by decreased expression of *Fth1*, and increased *de novo* unsaturated fatty acid biosynthesis. A similar observation was made in aged skeletal muscle: *Slc39a14* and *Hmox1* are upregulated, and the downregulation of *Tfr1* and *Fth1* expression contributes to the labile iron accumulation in skeletal muscle. A similar observation has been made in a macrophage-specific iron exporting protein, ferroportin (Fpn) ablation model*(4)*. In this model, monocyte-derived macrophages are indispensable in damaged skeletal muscle to secrete pro- or anti-inflammatory cytokines, which are necessary for the clearance of remnants and iron recycling. Although activated macrophages have a large portion of the intracellular labile iron pool, they have lower storage capacity and have to release iron into intracellular space to be utilized by newly formed myofibers*(45, 46)*. Macrophage-mediated iron recycling and muscular regeneration must be well-coordinated as macrophages provide a temporary storage site for iron to prevent oxidative damage and then subsequently supply nutrients for muscle regeneration.

Well-coordinated iron recycling and utilization between macrophages and newly formed myofibers has been proven to be critical to muscular regeneration. Thus, *Tfr1*-ablation in SCs results in an excessive amount of iron released from macrophages to the labile iron pool in skeletal muscle, but is not to be absorbed because of the defect in functional myofiber formation. Replenishment of Tfr1 protein via lentivirus infection partially decreases labile iron accumulation, prohibits fat biogenesis and promotes regeneration. The labile iron accumulation for one aspect is derived from myoglobin degradation by Hmox1 and additional NTBI *SC/KO* absorption is facilitated by Slc39a14, as the expression of both genes is upregulated in *Tfr1^SC/KO^* 5 dpi and remains at higher expression levels at 9 and 15 dpi, which definitely exacerbates iron accumulation and oxidative damage. It has been reported that in the absence of macrophage Fpn, iron sequestered inside the macrophages not only prevents muscle regeneration but also activates adipogenesis, leading to fat accumulation*(4)*. *Tfr1^SC/KO^* mice present *de novo* lipogenesis instead of fatty acid uptake *via Cd36* by upregulating expression of fatty acid synthase (*Fasn*) and the activation of the unsaturated fatty acid biogenesis pathway (e.g. *Scd1, Scd2, Fads1, Fads2*, and *Elvol5*). This is possible due to the activation of fibroadipogenic precursor cells (FAPs), a mesenchymal population located in the interstitial area of the skeletal muscle. FAPs are able to spontaneously differentiate into adipocyte or fibroblasts in *ex vivo* culture systems*(47)*. In *in vivo* system, FAPs are able to differentiate into adipocytes in degenerating dystrophic muscles while ectopic adipogenesis and fatty infiltration could be strongly inhibited by the presence of SC-derived myofibers*(48)*. Intramuscular fatty infiltration in skeletal muscle could be inhibited by IL-15 expression, possibly affecting FAP differentiation through Hedgehog signaling and other cytokines, e.g. IL-4 and IL-13 secreted from eosinophils to remove cellular debris, enhancing regeneration*(49–52)*. Skeletal muscle regeneration also requires network interactions among various cell types, such as endothelial cells, immune cells and motor neurons*(53)*. Numerous studies have demonstrated that muscle regeneration begins from 3 to 5 days after injury and peaks during the second week after injury*(54)*. Single-cell sequencing technology further demonstrates that skeletal muscle regeneration depends on a heterogeneous cell population, and regulated by various intra- and extracellular factors with the involvement of paracrine communication between SCs and nonmyogenic cells at different regeneration stages*(55, 56)*. In *Tfr1^SC/WT^* mice, we observed the peak of new myofiber formation at 5 dpi and almost complete of regeneration at 9 dpi as reported before*(56), Tfr1*-ablation fails to activate SCs and significantly delays the regeneration. In the TA of *Tfr1^SC/KO^* mice, both *Pax7* and *MyoD* mRNA were expressed at lower levels corresponding to decreased SC proliferation, and less newly generated myofibers at 5 dpi. We also did not observe any changes in the expression of ferroptosis related genes, such as *Gpx4* and *Ptgst2* at 5 dpi. However, as early as 5 dpi, some biomarkers that potentially contribute to the accumulation of labile iron, such as *Slc39a14* and *Hmox1* were upregulated in TA of *Tfr1^SC/KO^* mice, which could explain the phenomenon of massive iron accumulation at later time points. Even though *MyoD* expression was upregulated at 9 dpi (compared to *MyoD* expression at 5 dpi) in TA of *Tfr1^SC/KO^* mice, followed by the initiation of regeneration and a small number of eMyHC^+^ myofibers at small diameters, this may not have enough functional myofibers to utilize the iron and secrete cytokines to prevent adipogenesis. Skeletal muscle regeneration is further exacerbated by downregulation of Gpx4, a ROS scavenger, and Fth1, an important portion of ferritin, as well as upregulation of Slc39a14, an NTBI transporter and Ptgs2, involved in peroxidase generation at 9 dpi. The dysregulated iron homeostasis and adipogenesis results in the activation of ferroptosis. Ferritin degradation via reduced Fth1 expression is associated with the activation of ferroptosis, which is defined as ferritinophagy*(57, 58)*. We believe that *Tfr1*-ablation in SCs is a critical starting point of the skeletal muscle regeneration defect but not the ferroptosis marker, as reported before*(59)*. Most importantly, this *Tfr1* genetic deletion model and Tfr1-Slc39a14 functional switching recapitulate a physiological downregulation or dysfunction of Tfr1 in skeletal muscle and SCs during ageing in skeletal muscle of rodents. Ferroptosis activation is orchestrated by decreased Tfr1 membrane protein and Slc39a14 membrane enrichment to facilitate NTBI iron absorption contributing to labile iron accumulation in aged skeletal muscle. Moreover, FAPs derived abnormal fibrosis and adipogenesis in aged muscle contributes to abnormal lipid accumulation*(60)*, in together with iron accumulation and Fenton reaction leading to the activation of ferroptosis in skeletal muscle. These circumstances are also observed in aged skeletal muscle of rodents with the age-dependent risk of ferroptosis, which was reported elsewhere in the brain*(61, 62)*.

In additional to ferroptosis-mediated dysregulation of iron and lipid metabolism in skeletal muscle, we also observed systemic dysregulation of energy metabolism. Our electron transmission microscope images demonstrated that ferroptosis is followed by swollen mitochondria with irregular or disappeared cristae, immune cell infiltration, lysosome-mediated mitochondrial degradation or swollen mitochondria and lipid droplets in a membrane surrounded structure with lysosome. Along with the ferroptosis in the TA of *Tfr1^SC/KO^* mice, altered systemic energy metabolism is displayed with increased energy expenditure and expression of FGF21. FGF21 was initially discovered to be secreted from the liver regulating energy balance and glucose and lipid metabolism*(63)*. In skeletal muscle under healthy and physiological conditions, its expression remains at a lower level. However, FGF21-induced muscle atrophy/weakness during fasting or FGF21 overexpression *in vivo* in muscle is sufficient to induce autophagy and muscle loss by 15%*(64)*. Its deletion could protect against muscle loss and weakness during fasting, which is accompanied by a significant reduction of mitophagy flux*(64)*. Other than its pathological function in skeletal muscle, FGF21 is a potent stimulator of adipocyte thermogenesis and nutrient metabolism to improve insulin sensitivity and reduce hepatic lipid accumulation*(63, 65)*. Studies have suggested that autocrine and paracrine actions of FGF21 are able to induce thermogenic effects in adipocytes to improve glucose metabolism, lipid profiles and anti-obesity effects*(66–68)*. Between *Tfr1^SC/WT^* and *Tfr1^SC/KO^* mice, accompanied by increased EE but no significant difference in ratio of VCO_2_/VO_2_ and the adipocyte thermogenic signaling pathway was not observed, indicating that a Ucp1-independent mechanism may be involved. Ucp1-independent thermogenic pathways, such as creatine metabolism, calcium cycling, and amino acid uncoupling, promote systemic energy metabolism*(69–71)*. FGF21 or FGF21 mimetic antibody stimulates brown or white adipocyte thermogenesis in a Ucp1-independent manner, which may act via directly promoting the host metabolic activity*(72)* or the FGFR1/bKlotho complex*(73)*, respectively.

Meanwhile, resident and monocyte-derived macrophages also contribute to the skeletal muscle regeneration at different stages. The depletion of macrophage in Tg-ITGAM-DTR mice impairs regeneration and results in the lipid accumulation in the skeletal muscle*(74)*. Macrophages in skeletal muscle play a fundamental role in muscle repair and debris clearance. Upon initiation of muscle damage, infiltrated monocytes/neutrophil in injured areas differentiate into proinflammatory macrophage (M1 Macrophage) with exposure to interferon-(IFN) γ and tumor necrosis factor (TNF)α to phagocyte necrotic muscle debris *(75)*. M2 macrophage polarization majorly presents at the advanced stage of tissue repair and wound healing in concert with the secretion of IL-14 and IL13 from Th2 cytokines. Alternatively, M2 macrophages are also associated with the fibrosis in *mdx* mouse model, indicating M2 macrophages may have alternative function under pathological condition *(76, 77)*. Despite the compelling evidence of different macrophage subtypes during muscle repair, we identified a significant accumulation of M2 macrophages in the TA of *Tfr1^SC/KO^* mice, leading to the development of fibrosis and unexpected macrophage-derived foam cell differentiation. Form cells are associated with the development of atherosclerosis, and are also implicated in proinflammatory cytokine secretion, different inflammatory cells recruitment and fibrotic collagen accumulation, which further exacerbates tissue function and impairs tissue repair*(78)*. However, the pathological function of foam cells in skeletal muscle requires further exploration.

In summary, the current investigation reveals that SCs with ablation of *Tfr1* impairs skeletal muscle regeneration with the activation of ferroptosis. This process is accompanied by Tfr1-Slc39a14 functional switch to mediate NHI absorption and labile iron accumulation in the skeletal muscle. This phenomenon is recapitulated in the skeletal muscle of aged rodents, which may shed light on the development of anti-ageing strategies.

## Materials and Methods

### Animals and treatment

*C57BL/6J* mice were purchased from the Center of Guangdong Experimental Animal Laboratory and housed in a temperature and humidity controlled and ventilated specific pathogen free (SPF) cages at animal facility of Guangdong Institute of Microbiology. All animal handling and procedures were approved by the Animal Care and Use Committee at Guangdong Institute of Microbiology [Permission #: GT-IACUC201704071]. All experimental mice were placed on a 12-hr light: dark cycle with *ad libitum* access to food and water.

Mice with *Tfr1*-spepcifc deletion in SCs were generated by crossing mice carrying *Pax7-CreER* and *Tfr1^fl/fl^* allele. The genotype was *Pax7-CreER*;*Tfr1^fl/fl^* designated as homozygous (*Tfr1^SC/KO^*)and as control littermate (*Tfr1^SC/WT^*). SCs-specific *Tfr1* deletion was generated by intraperitoneal (*i.p*.)-injection of Tamoxifen (T5648, Sigma) dissolved in corn oil for 7 consecutive days at dose of 15 mg/ml as described before*(24)*. *Pax7-CreER* mice were shared by Dr. Dahai Zhu from Institute of Basic Medical Sciences (Chinese Academy of Medical Sciences), originally purchased from Jackson Laboratory (Stork No: 017763) and *Tfr1^fl/fl^* mice were directly purchased from Jackson Laboratory (Stok No: 028363).

To induce muscle injury, Cardiotoxin (CTX, 0.5 nmol, 100 μl) was intramuscularly administrate. For the drug treatment, CTX-injured mice were *i.p*.-injected with Saline or Ferrostatin-1 (2 μmol/kg, SML0583, Sigma), a ferroptosis inhibitor for 30 days. For lentivirus administration, mice with CTX-injured TA muscle were intramuscularly administrated with control or lentivirus with Tfr1-expression.

### Lentivirus packaging

293T cells seeded in 10 cm plate at 95% confluency were transiently transfected with shuttling vector (pCDH-Tfr1, kindly shared by Dr. Fudi Wang or empty vector) and packaging vectors (pVSVg and psPax2) by TransIT X2 (MIR60000, MirusBio). 12 hrs after transfection, culture medium was replaced with fresh DMEM supplemented with 10% fetal bovine serum (FBS,01010102, Trinity). Medium containing lentivirus were harvested respectively at 36 and 60 hrs after transfection. Lenti-X concentrator (PT4421-2, Clontech) was mixed at the ratio of 1:3 and incubated at 4 °C for a short time. The mixture was then centrifuged to obtain a high titer virus containing pellet, which was resuspended in 500 μl PBS/saline solution and stored in −80 °C freezer.

### RNA isolation and real-time PCR

Total RNA of skeletal muscles, liver, iBAT, iWAT or eWAT was extracted with TRIzol™ reagent (1596018, Thermo Fisher) and reverse transcribed by utilizing 5x All-In-One Master Mix (G485, AbmGood) according to the manufacture’s instruction. cDNA was used to analyze gene expression by Power SYBR Green Master Mix (4367659, Thermo Fisher) on a QuantStudio 6 Flex Real-Time PCR System (Thermo Fisher). The primer sequences for qRT-PCR were listed in Table 1.

**Table 1.**
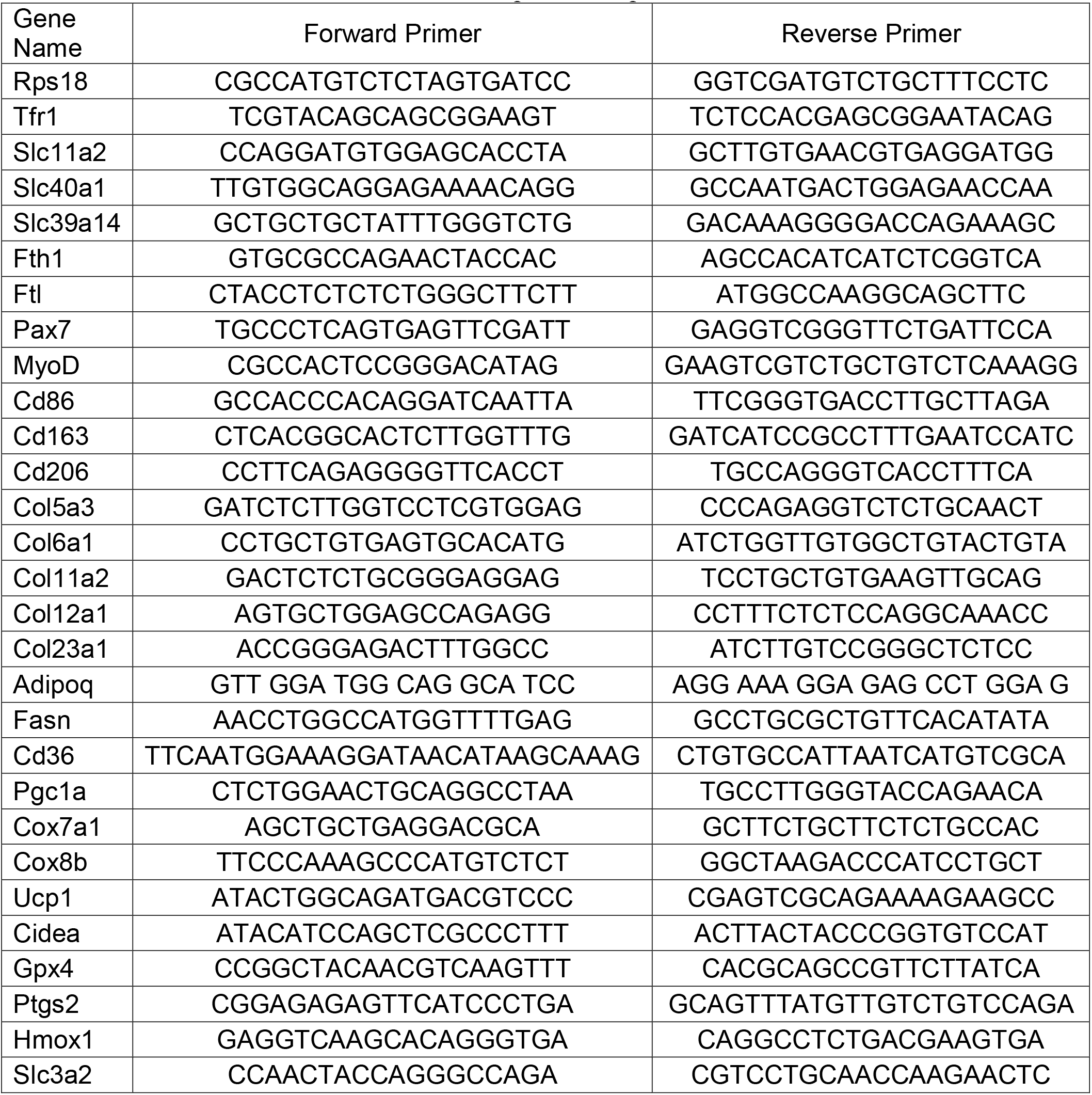
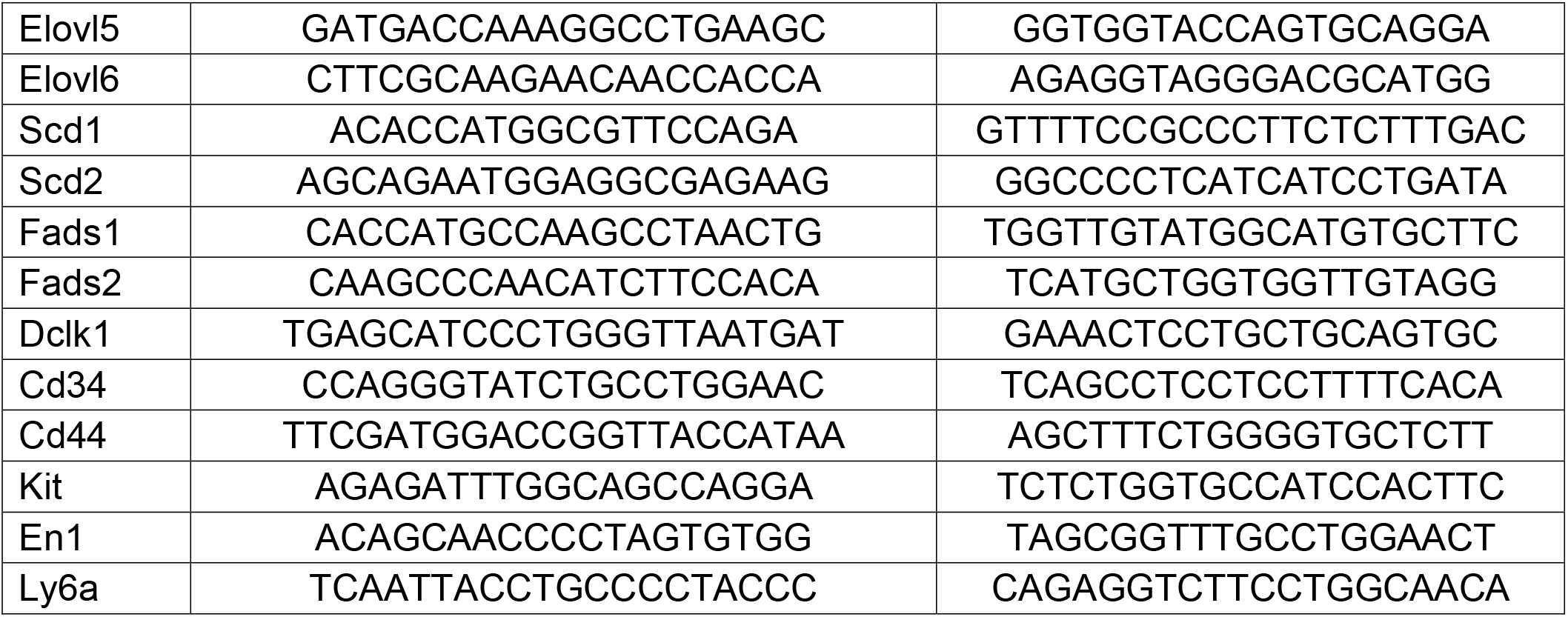
Primer sequence for qPCR.

### RNAseq and bioinformatic analysis

Total RNA samples from *C57BL6/J* mice at different age (n=3 per group) or *Tfr1^SC/WT^* and *Tfr1^SC/KO^* (n=5 per group) before or after CTX-inury at 30 dpi were sequenced using a BGI-SEQ2000 platform (Beijing Genomics Institute). Raw RNA-seq reads in FASTQ format were quality checked with FASTQC algorithm, and low-quality reads were trimmed using the FASTX-Toolkit. High-quality reads were aligned to the mouse genome (GRCm38/mm10) using HISAT2*(79)* and assembled against mouse mRNA annotation using HTSeq*(80)*. Differentially expressed genes (DEGs) were analyzed by using DESeq2 package in R*(81)*. Genes were considered to be significantly upregulated or downregulated at *padj*<0.05. Heatmaps were generated using the pheatmap package in R based on raw count of DEGs. Gene ontology (GO) analysis was performed using the R package, named clusterProfiler for DEGs (either up- or down-regulated)*(82)*. DEGs (*padj*<0.05) were further analyzed using Gene Set Enrichment Analysis (GSEA)*(83)*. Both upregulated and downregulated genes were functionally categorized with the GO and Hallmark gene sets.

### Iron assay

Skeletal muscles (TA, EDL, Sol and Gas) non-heme iron levels or serum iron levels were determined following a standard protocol as described before*(84)*. In short, weight skeletal muscles were homogenized in H_2_O and equal volume of acid solution (10% trichloroacetic acid in 3 M HCl) added. Samples were digested for 1-hr at 100 °C. 75 μl digested sample or iron standard was mixed with 75 μl ferrozine solution (1mM Ferrozine, 3M Sodium Acetate and 1% mercaptoacetic acid), followed with incubation at 37 °C for 1-hr before the colorimetric was read at 565 nm by a microplate photometer (Thermo Fisher). The iron level of each sample was normalized by the weight of skeletal muscles and presented as micrograms of iron per gram of wet tissue weight.

### Oil Red O staining

4% PFA fixed TA sections were stained in an Oil Red O (ORO) solution following a standard protocol as described before. The nuclei were counter-stained with haematoxylin before mounted with glycerol-containing mounting medium.

### Perls’ Prussian’s Blue staining

Non-heme iron staining was performed by utilizing a standard Perl’s Prussian Blue Staining protocol as previously described*(84)*. Images were visualized and captured with a light microscope.

### Masson’s Trichrome stain

Masson’s Trichrome stain was performed using Masson’s Trichrome Stain Kit (Aniline Blue) following the manufacturer’s instructions (MA0123, Meilunbio). Briefly, Masson’s Trichrome stain was performed on 10 μm cryosection of TA muscle fixed with 95% alcohol for 20-min. Sections then were incubated different solutions supplemented in Masson’s Trichrome Stain Kit. At the end, the section was dehydrated with 95% alcohol for 10s, two rinses in anhydrous alcohol for 10 s and 2 rinses in xylene for 1 min each. The sections were mounted with Neutral balsam for imaging and fibrosis quantification. TA fibrosis quantification was performed by using Image J. CVF (Collagen volume fraction), which is calculated to be the ratio of the collagen-positive blue area versus the total tissue area.

### Hematoxylin and Eosin (H&E) staining

TA sections at 10 μm were stained with hematoxylin and eosin solution by following a standard protocol as described before*(44)*. Sections were dehydrated and mounted with DPX Mountant (44581, Sigma). Histological images were visualized and captured by a light microscope.

### Protein isolation and western blot

Total protein lysates were prepared and resolved on SDS-PAGE as described before*(85)*. Protein band on a PVDF membrane was probed with primary antibodies (Cav1: D161423, Tubulin: D225847, Sangon Biotech; Tfr1: ab84036, Mitochondrial Complex: ab110413, Abcam; PGC1α: ab3242, Millipore;, Pparγ: sc-7273, Fth1: sc-376594, Ftl: sc-74513, Santa Cruz Biotechnology; Gpx4: A1933, Abclonal; Slc39a14: PA5-21077, Thermo Fisher; Acc:3676; Nrf2: 12721, Cell Signaling Technology) overnight, followed with secondary antibodies incubation at RT for 1-hr. Images were acquired using the ChemiDocTM Imaging System (Bio-Rad).

### Myofiber isolation, culture and immunofluorescence staining

Single myofibers were isolated from extensor digitorum longus (EDL) muscle following the method as described before*(24, 86)*. Briefly, EDL muscle was isolated and incubated in digestion medium containing 0.2% collagenase for 75-min. After digestion, myofibers were transferred to a horse serum coated 24-well plate and gently washed for three times with the washing medium (DMEM supplemented with 10% FBS: fetal bovine serum and 1% P.S.: Penicillin and Streptavidin). For non-cultured myofiber, at least 100 myofibers for each group were fixed with 4% paraformaldehyde (PFA, P6148, Sigma) and immunostaining was performed following a standard protocol. For myofiber culture, single myofiber was either cultured in a horse-serum-coated 24-well plate (for non-attached culture) or a collagen-coated 24-well plate (for attached culture). In each well, ~20 myofibers were washed for three times before replaced with culture medium (DMSM supplemented with 20% FBS, 1% P.S. and 1% CEE: chicken embryo extract, (C19041654, USBiological) with or without 4-OH-Tamoxifen (1 μm, H7904, Sigma). For non-attached culture, after 72-hr culture, single myofibers with SCs-cluster were fixed with 4% PFA and immunostaining was performed following a standard protocol. For attached culture, after 72-hr culture, SCs cluster attached the culture plate and proliferated for 4-6 days until reaching 85% confluency, followed with 3 days differentiation with 2% horse-serum. The myotubes were fixed with 4% PFA, followed with immunostaining. Single myofiber or differentiated myotubes were permeabilized with 0.5% Triton-100 for 10 mins before blocking with sterilized PBS containing 3% BSA, 5% goat serum and 0.5% tween 20). Primary antibodies (DSHB: Pax7: PAX7, eMyHC: F1.652, MHC:20-s, Abcam: Tfr1: ab84036 and Active Motif: MyoD:39991; Ki67:ZM-0167, ZSGB-Bio) were incubated overnight at 4 °C and secondary antibodies at RT for 1 hr at dark. Myofibers were mounted with DAPI-containing mounting medium (F6057, Sigma). To quantify the number of SCs, Pax7^+^ SCs on myofibers with Tfr1, MyoD, and Ki67 expression were counted. Images were visualized and captured with EVOS Cell Imaging Systems (EVOS FL, Thermo Fisher) or Confocal Microscope (Zessie 710).

### Myoblast isolation and culture

Primary myoblasts were isolated from 2-week old *Tfr1^fl/fl^* mice as described before*(24)*. Primary myoblasts were cultured on a collagen-coated tissue cell culture dish in Nutrient Mixture F-10 Ham (N6635, Sigma) supplemented with 20% FBS, 1% P/S, 5ug/L FGF-basic (100-18B, PeproTech). For proliferation analysis, primary myoblasts were infected with adenovirus expressing Cre recombinase at 50 MOI to induce Tfr1-deletion, followed by incubation with EdU for 24 hrs before harvest for immunostaining by utilizing the Click-iT™ EdU Cell Prolieration Kit (C10337, Thermo Fisher).

### Cryosection and immunofluorescence staining

TA muscle was dissected, mounted, frozen and sectioned at 10 μm as described before *(24)*. TA section was fixed with 1% PFA, and antigen retrieval was performed with Tris-EDTA buffer (Tris 1.21g and EDTA 0.37g dissolved in 1L ddH_2_O, pH 9.0) for 1-hr at 100 °C. Sections were permeabilized with 0.5% Triton-100 for 10-min and blocked with blocking buffer (PBS with 3% BSA and 5% goat serum) for 1 hr. Primary antibodies was incubated O.N. at 4°C (Wheat Germ Agglutinin (WGA):W32466, Thermo Fisher; LaminB2: 05-206, Millipore; DSHB: Pax7: PAX7, type I: #BA-D5, type IIA: SC-71, type IIB: BF-F3 and type IIX: 6H1-s, Perilipin: 9349, Cell Signaling Technology), followed by secondary antibodies incubation at R.T. for 1-hr in dark room. Nuclei were counterstained with DAPI-containing mounting medium (F6057, Sigma). The image was visualized and captured with the EVOS Cell Imaging Systems (Thermo Fisher).

### Transmission electron microscopy

TA muscle injury was induced by intramuscular injection of CTX for both *Tfr1^SC/KO^* and control littermates. 15 days after injury, TA samples (1 mm x 1 mm x 1mm) were quickly dissected and immediately fixed in 4% phosphate-glutaraldehyde. Each sample was dehydrated, permeabilized, embedded, sectioned at 60-80 nm and mounted. For each sample, five fields of view were randomly selected and the images were captured.

### Treadmill exhaustion test

Treadmill exhaustion test was performed for both *Tfr1^SC/KO^* and control littermates before and after CTX-induced muscle regeneration. The treadmill running protocol was started with an adaptation period of 10 m/min for 20-min before an increase of 2 m/min every 20-min until fatigue response initiated. The treadmill running protocol was terminated when mice no longer responded to 5 consecutive fatigue stimuli. Upon fatigue initiated, mice were quickly removed from treadmill running lane. Treadmill running time and distance was recorded and calculated for all mice.

### Flow cytometry

Single-cell suspensions were incubated with purified anti-CD16/CD32 Abs (clone 2.4G2, Sungene Biotech, Tianjin, China) for 15 min to block Fc receptors. After wash, cells were stained with eFluor 450-anti-mouse CD45 (clone 30-F11, Invitrogen), Percp-Cy5.5-anti-mouse/human CD11b (clone M1/70, Biolegend), PE-Cy7-anti-mouse F4/80 (clone BM8, Biolegend), APC-anti-mouse CD86 (clone GL-1, Biolegend), FITC-anti-mouse CD206 (clone C068C2, Biolegend) or isotype controls at 4□ for 15 min and detected by flow cytometry (FACSVerse, BD). Data were analyzed using FlowJo software (V10). Macrophages were identified as CD45^+^/CD11b^+^/F4/80^+^, and the percentage of pro-inflammatory (CD86^+^) and antiinflammatory (CD206^+^) macrophages were analyzed and shown.

### Statistical analysis

Experiment results were presented as mean ± SEM. Bar plots and statistical analysis were generated using GraphPad Prism 7 by using Unpaired Student’s T tests or One-Way ANOVA with *p*< 0.05 were considered significantly different. Representation of the *p-values* was as follows: *p < 0.05, **p < 0.01, ***p≤0.001, N.S.: not significant (*p*≥0.05).

## Author Contributions

LW. X., Y.L. and X. Z. designed the experiment. HR. D., SJ. C., XP. Y., and XS. D. developed and optimized the methodology. HR. D., Z. L., XD. M., Y. T., SS. Z., BD. L., GC. C., ZC. Y., XP. Yao, L. Y., XY. C., J. S., H. C., YL. Y., GH. X., HJ. L., WD. W. Z. C., and JC. L collected and analyzed the data. GH. P., L. G. and ML. H. raised the experimental mice. LW. X acquired the grants and drafted the manuscript. We appreciated Mr Guanshen Liu from Biomarker Technologies Corporation for assisting us the RNA sequencing and Ms Xiaochen Wang for the figure editing.

## Data and materials availability

RNAseq data are available upon request.

## Supplementary Materials

**Figure S1.**
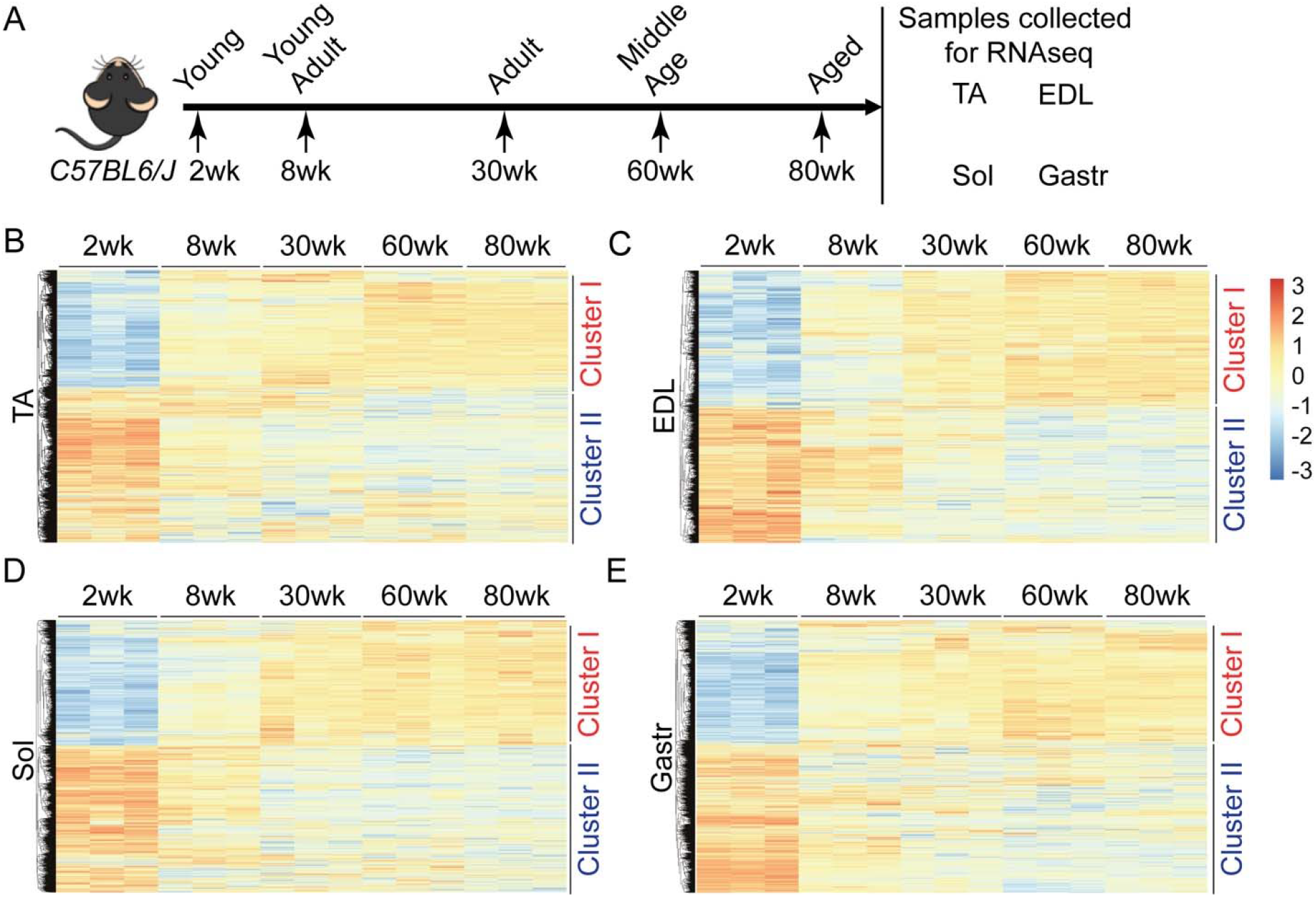
Related to Figure 1 Gene expression profile in four skeletal muscles (TA, EDL, Sol and Gas) across five different age stages (2wk-, 8wk-, 30wk-, 60wk- and 80wk-old). (A) Scheme of experimental design; (B) Heatmap of gene expression in TA from *C57BL/6J* mice across five different age-stage (n=3/group); (C) Heatmap of gene expression in EDL from *C57BL/6J* mice across five different age-stage (n=3/group); (D) Heatmap of gene expression in Sol from *C57BL/6J* mice across five different age-stage (n=3/group); (E) Heatmap of gene expression in Gas from *C57BL/6J* mice across five different age-stage (n=3/group).

**Figure S2.**
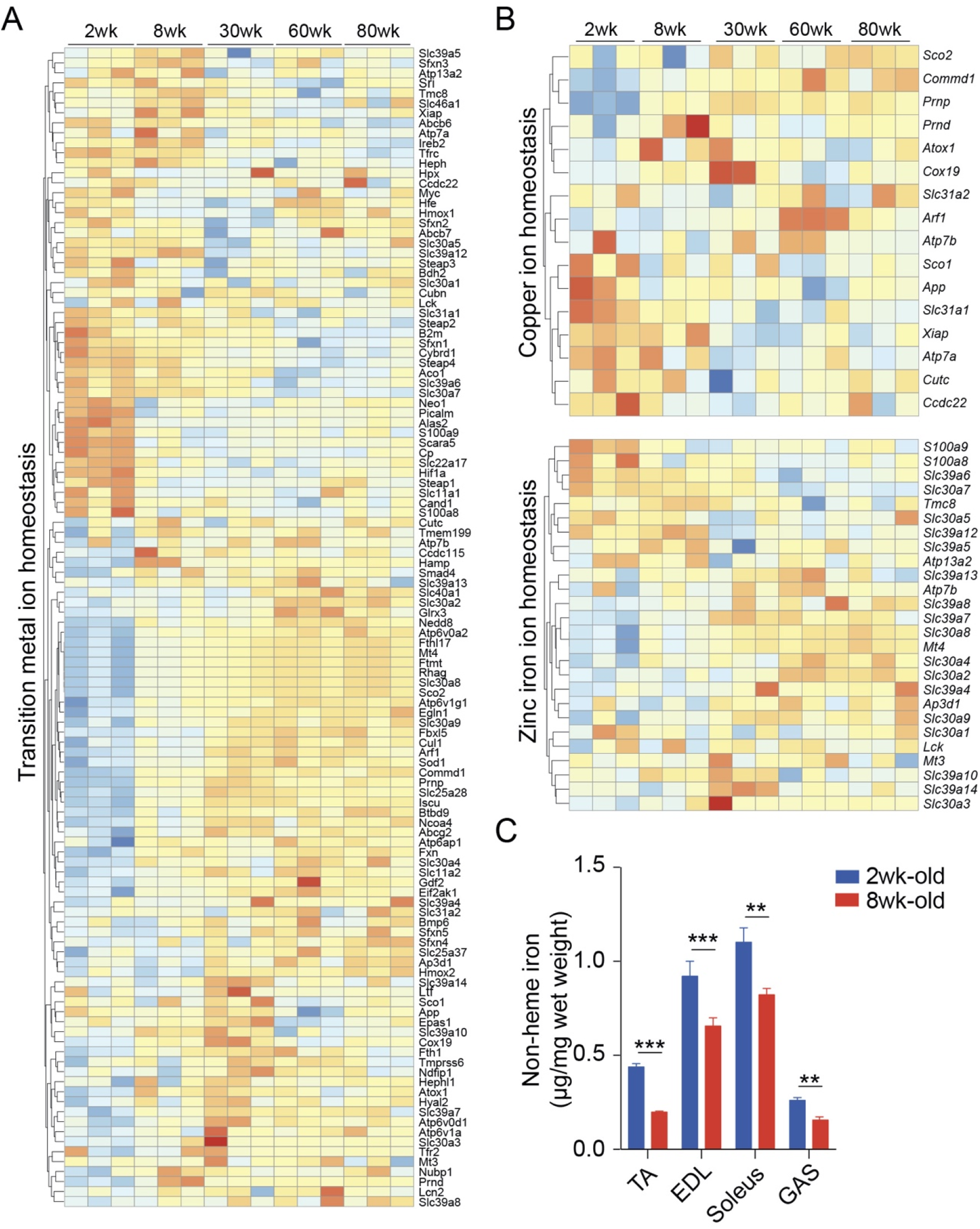
Related to Figure 1 Gene expression profile in four skeletal muscles (TA, EDL, Sol and Gas) across five different age stages (2wk-, 8wk-, 30wk-, 60wk- and 80wk-old). (A) Heatmap of transition metal ion homeostasis related gene expression profile in TA muscle across five different age stages (n=3/group); (B) Heatmap of copper and zinc ion homeostasis related gene expression profile in TA muscle across five different age stages (n=3/group); (C) Total non-heme iron levels in four skeletal muscles between 2wk- and 8wk-old *C57BL/6J* mice (n=6/group). N.S.: not significant, **P < 0.01, ***P < 0.005, by 2-sided Student’s t-test. Data represent the mean ± SEM.

**Figure S3.**
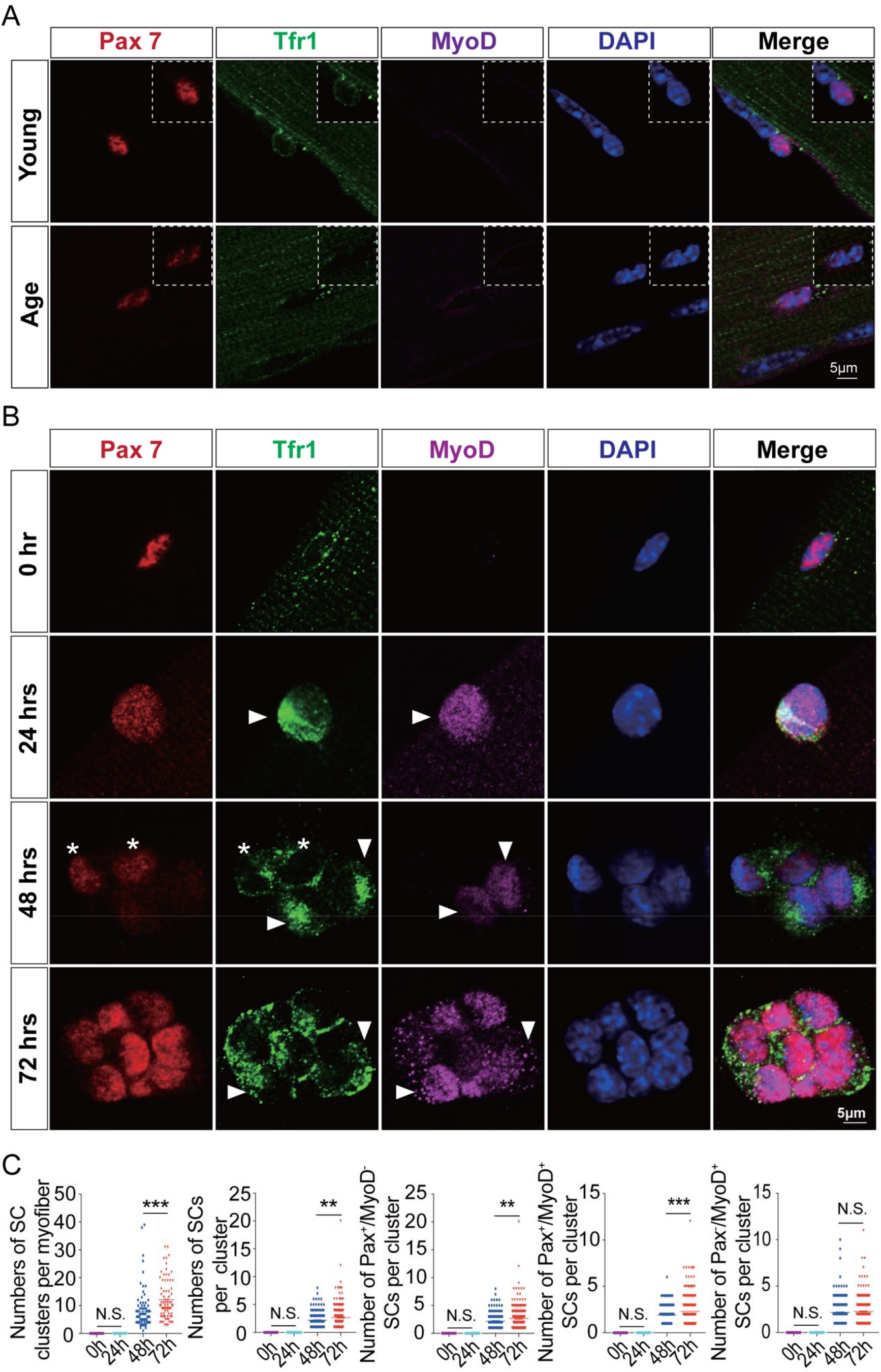
Related to Figure 1 Tfr1 protein presents in proliferative SCs. (A) Representative images of myofibers from 8wk- and 80wk-old *C57BL/6J* mice (n>50 myofibers from 5 mice/group). Immunofluorescence of Pax7 (red), Tfr1 (green), MyoD (pink) and DAPI (blue) staining revealed Tfr1 protein is lowly expressed in SCs of old *C57BL/6J* mice; (B) Representative images of SCs cluster on single myofiber from adult *C57BL/6J* mice culture for different length of time (0, 24, 48 and 72 hrs post single myofiber isolation); (C) Number of SC clusters, Pax7^+^ SCs per cluster, and Pax7^+^/MyoD^−^, Pax7^+^/MyoD^+^, and Pax7^−^/MyoD^+^ SCs per SCs cluster (n>50 myofibers). N.S.: not significant, **P < 0.01, ***P < 0.005, by 2-sided Student’s t-test. Data represent the mean ± SEM.

**Figure S4.**
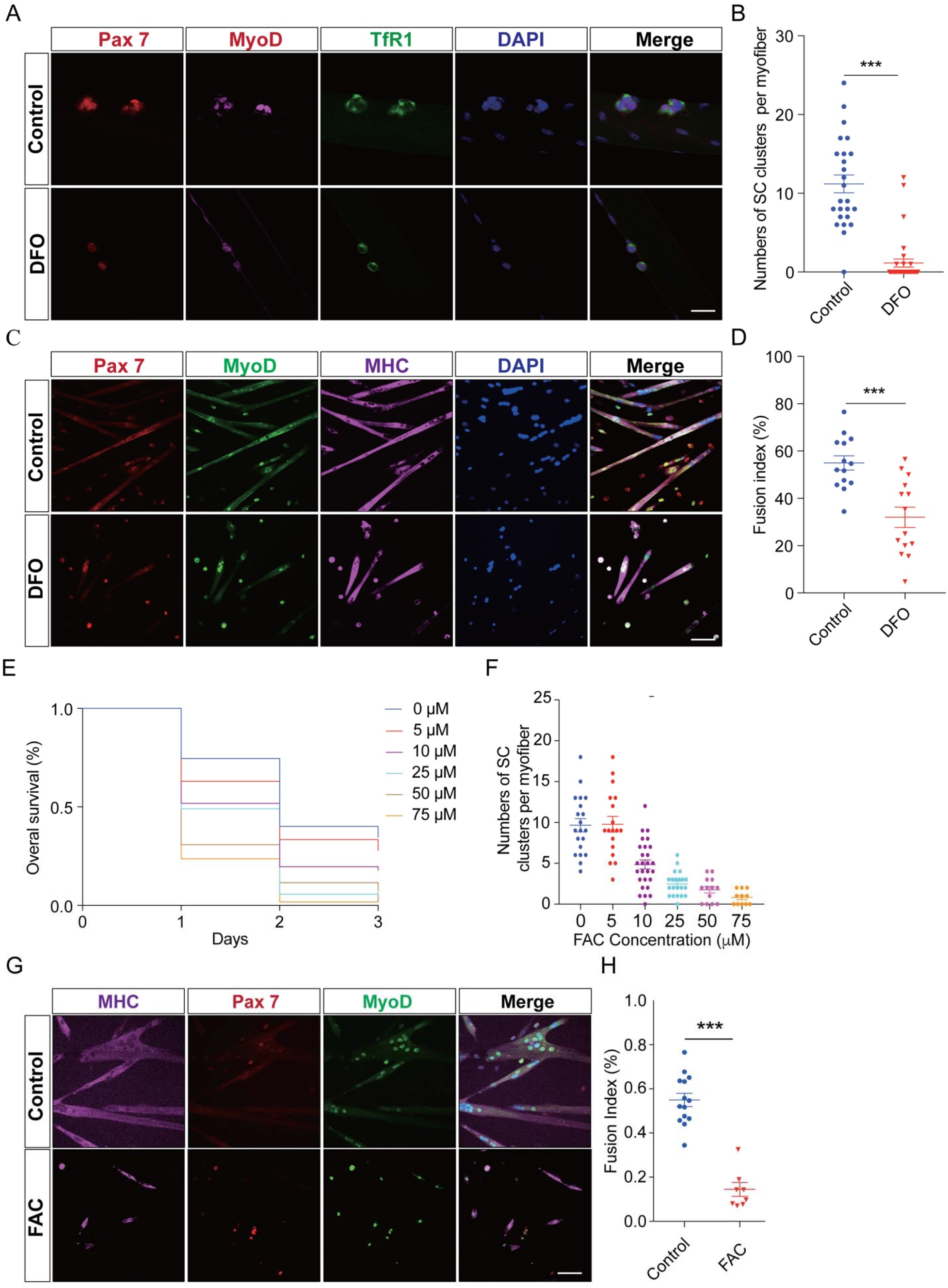
Related to Figure 1 Dysregulation of iron homeostasis inhibits SCs proliferation and differentiation. (A) Representative images of myofibers from adult *C57BL/6J* mice treated with DFO (an iron chelator) or vehicle (n>50 myofibers from 5 mice/group). Immunofluorescence of Pax7 (red), Tfr1 (green), MyoD (pink) and DAPI (blue) staining revealed iron deprivation inhibit SCs proliferation; (B) Number of SC clusters per myofiber; (C) Representative images of SCs differentiated into myotube with DFO treatment or vehicle. Immunofluorescence of Pax7 (red), MyoD (green), MHC (pink) and DAPI (blue) staining revealed iron deprivation inhibits SCs differentiation; (D) Summary of fusion index between control and DFO treatment group; (E) Overall survival rate of *ex vivo* cultured myofibers treated with different concentration of FAC (0, 5, 10, 25, 50, and 75 μm). The number of survived myofiber was counted every 24 hrs. (F) Number of SC clusters per myofiber treated with different concentration of FAC (0, 5, 10, 25, 50, and 75 μm) and cultured for 72 hrs; (G) Representative images of SCs differentiated into myotube with FAC treatment (25 μm). Immunofluorescence of Pax7 (red), MyoD (green), MHC (pink) and DAPI (blue) staining revealed FAC treatment at higher concentration inhibits SCs differentiation. N.S.: not significant, ***P < 0.005, by 2-sided Student’s t-test. Data represent the mean ± SEM.

**Figure S5.**
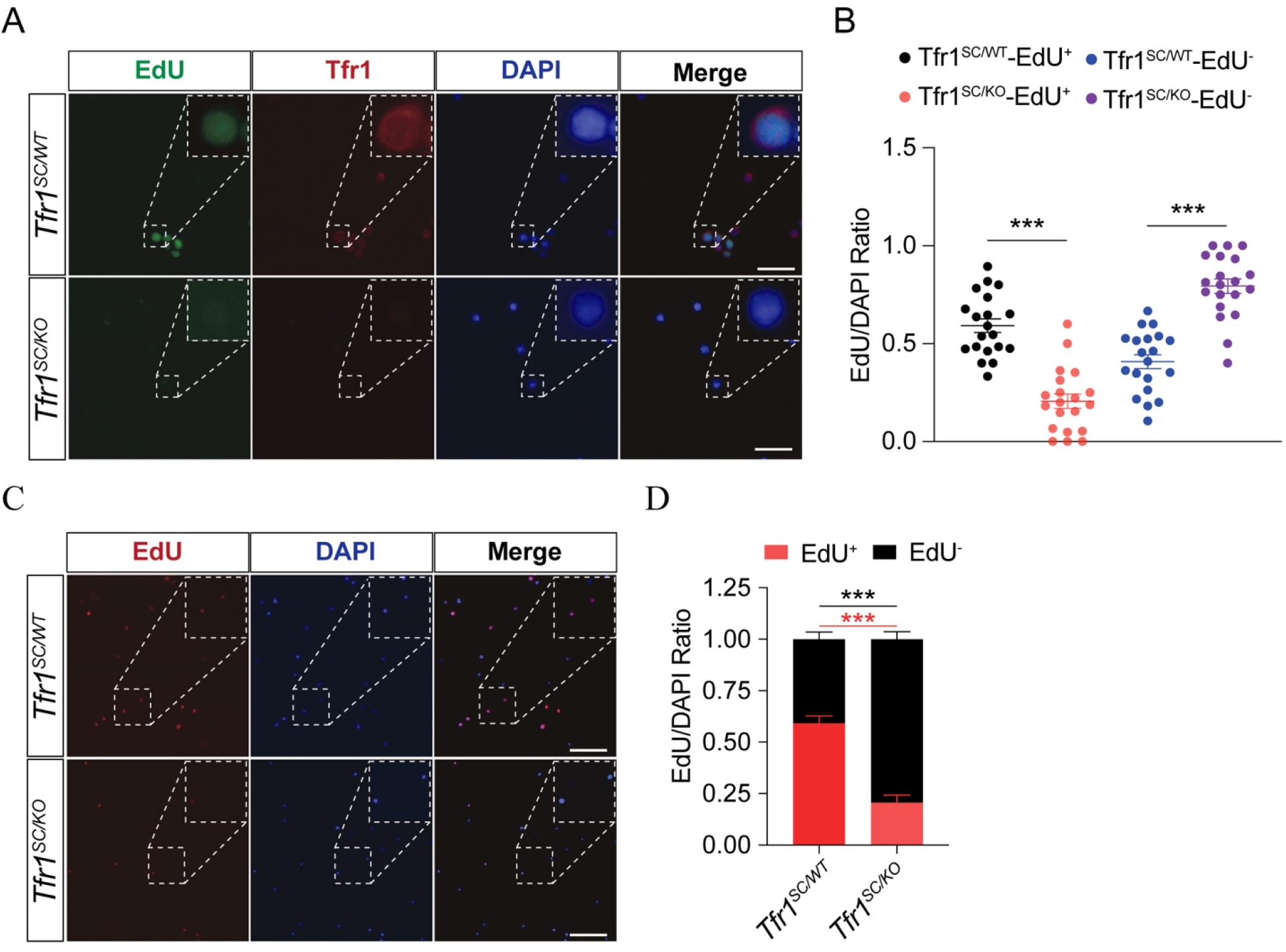
Related to Figure 2 *Tfr1*-ablation in myoblasts prevents the proliferation. Pax7^+^ SCs were isolated from *Tfr1^fl/fl^* mice and culture with F10 medium. Myoblasts were treated with Adenovirus expression Cre recombinase or GFP as control. 24 hrs later, myoblasts were further treated with EdU for 24 hrs before immunostaining. (A) Representative images of myoblasts immunostaining with EdU (green), Tfr1 (red) and DAPI (blue) indicates that Tfr1-ablation inhibits myoblast proliferation; (B) Number of EdU^-^ and EdU^+^ myoblasts in total DAPI^+^ myoblast from both control and Tfr1-delted group; (C) Representative images of myoblasts immunostaining with EdU (red) and DAPI (blue) indicates that *Tfr1*-ablation inhibits myoblast proliferation; (D) Stacking bar graph showing the ratio of EdU^-^ and EdU^+^ myoblasts. N.S.: not significant, ***P < 0.005, by 2-sided Student’s t-test. Data represent the mean ± SEM.

**Figure S6.**
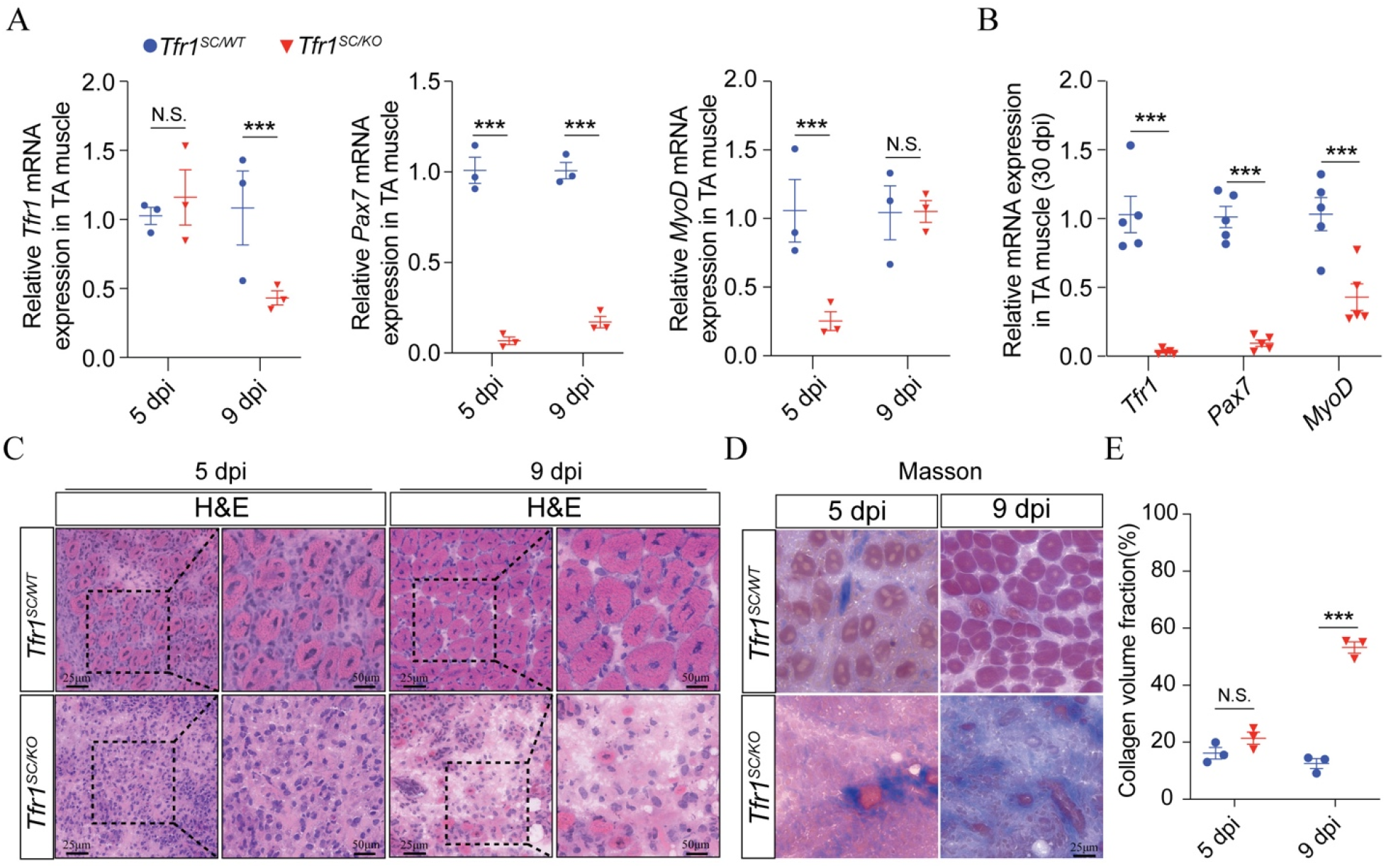
Related to Figure 3 *Tfr1*-ablation in SCs inhibits SCs proliferation and skeletal muscle regeneration. (A) qPCR analysis of Tfr1, Pax7 and MyoD expression in TA muscle from *Tfr1^SC/WT^* and *Tfr1^SC/KO^* mice after CTX-induced injury at 5 and 9 dpi; (B) qPCR analysis of *Tfr1, Pax7* and *MyoD* expression in TA muscle of *Tfr1^SC/WT^* and *Tfr1^SC/KO^* mice after CTX-induced injury at 30 dpi; (C) Representative images of TA section H.E. staining of *Tfr1^SC/WT^* and *Tfr1^SC/KO^* mice after CTX-induced injury at 5 and 9 dpi; (D) Representative images of TA section Masson staining of *Tfr1^SC/WT^* and *Tfr1^SC/KO^* mice after CTX-induced injury at 5 and 9 dpi; (E) Summary of collagen volume fraction on TA section of *Tfr1^SC/WT^* and *Tfr1^SC/KO^* mice after CTX-induced injury at 5 and 9 dpi. N.S.: not significant, ***P < 0.005, by 2-sided Student’s t-test. Data represent the mean ± SEM.

**Figure S7.**
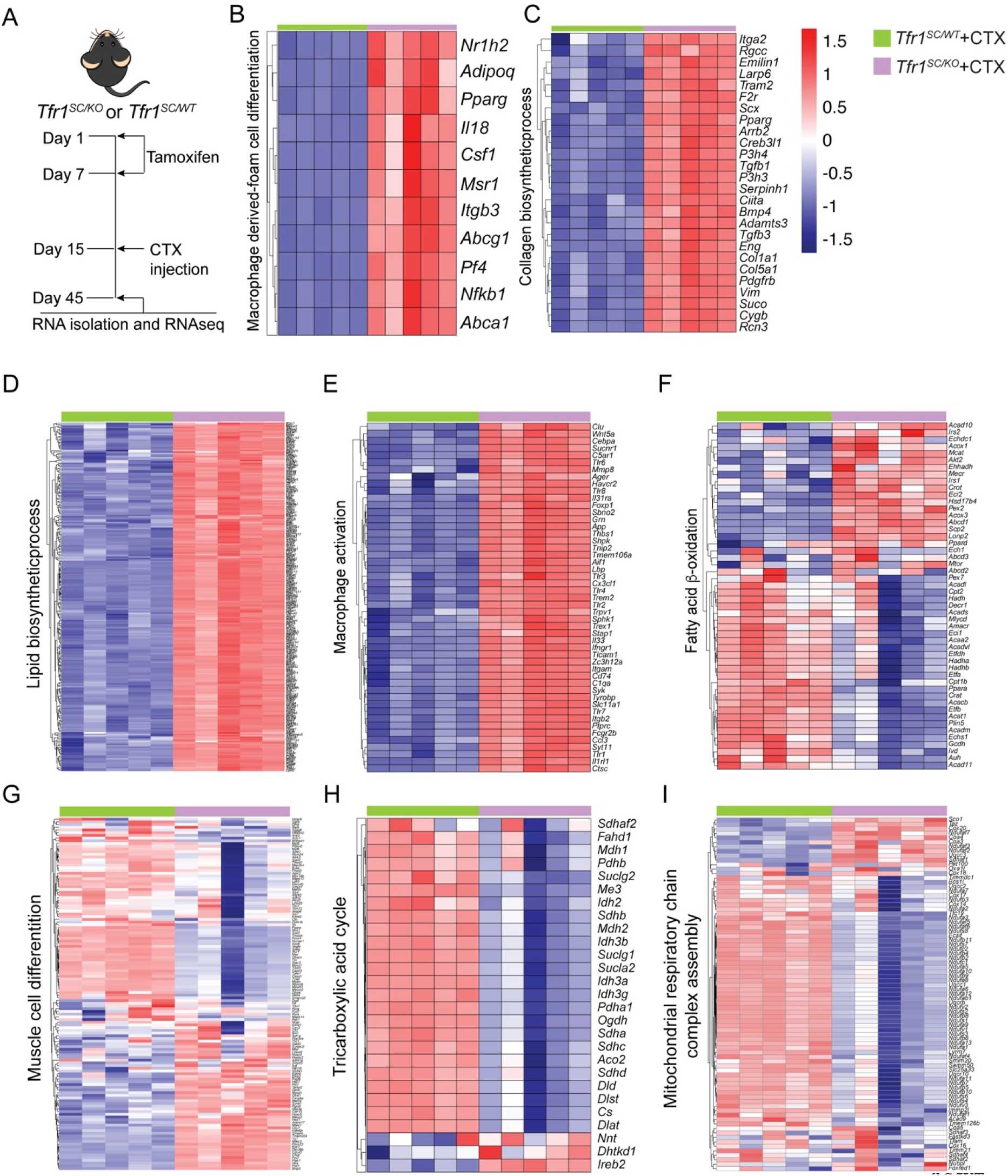
Related to Figure 4 Gene expression profile in TA muscle between *Tfr1* and *Tfr1^SC/KO^* mice after CTX-induced injury at 30 dpi. (A) Timeline for tamoxifen-induced *Tfr1* deletion and CTX-induced TA muscle injury; (B) Heatmap of macrophage-derived foam cell differentiation related gene expression; (C) Heatmap of collagen biosynthetic process related gene expression; (D) Heatmap of lipid biosynthetic process related gene expression; (E) Heatmap of macrophage activation related gene expression; (F) Heatmap of fatty acid β-oxidation related gene expression; (G) Heatmap of muscle cell differentiation related gene expression; (H) Heatmap of tricarboxylic acid cycle related gene expression; (I) Heatmap of mitochondrial respiratory chain complex assembly related gene expression.

**Figure S8.**
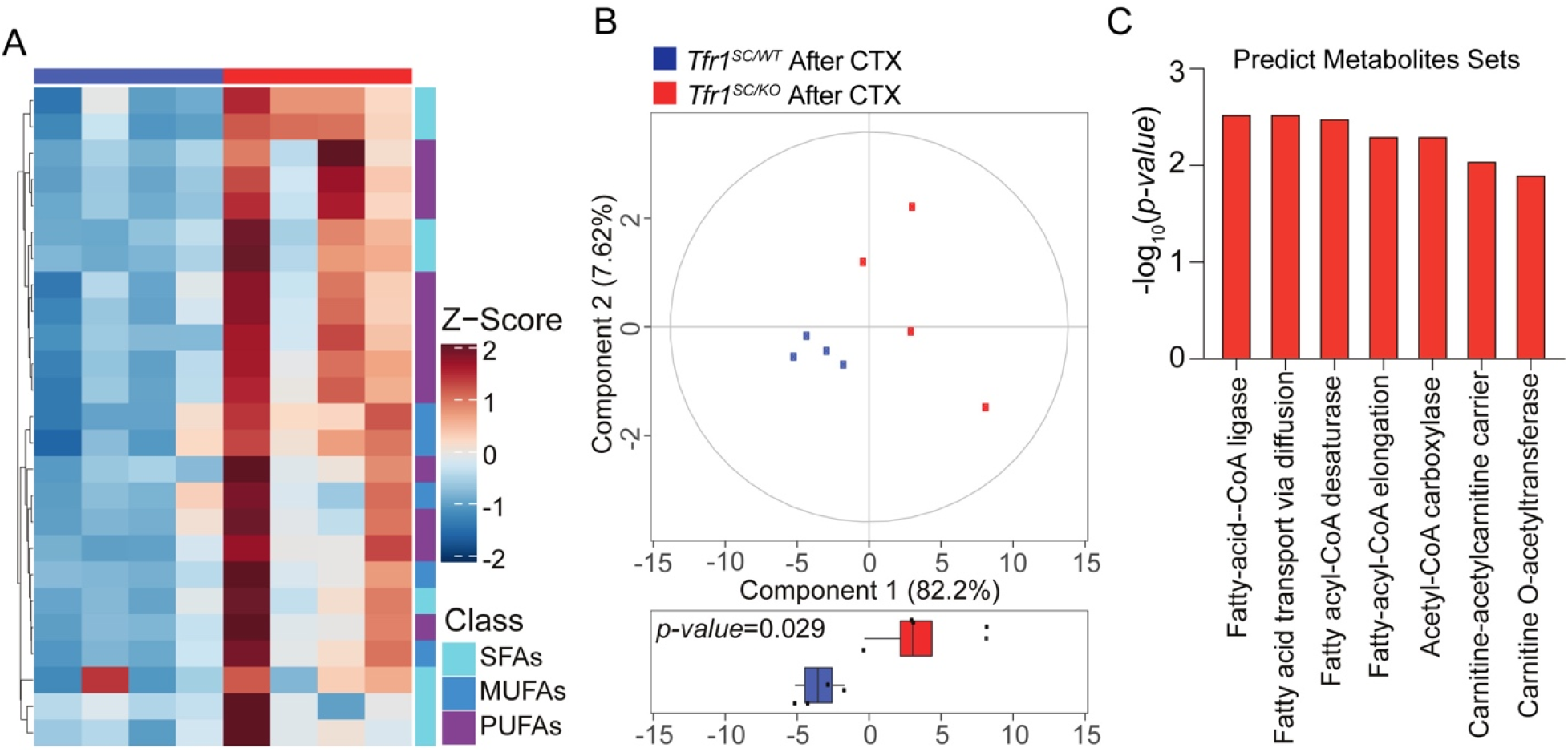
Related to Figure 5 TA of *Tfr1^SC/KO^* mice has different lipogenesis profiling. (A) Heatmap presenting the amount of SFAs, MUFAs and PUFA between *Tfr1^SC/WT^* and *Tfr1^SC/KO^* mice after CTX-induced injury at 15 30 dpi; (B) PCA of lipid profiling between *Tfr1^SC/WT^* and *Tfr1^SC/KO^* mice after CTX-induced injury at 15 30 dpi; (C) Bar graph showing the predicted pathway enrichment.

**Figure S9.**
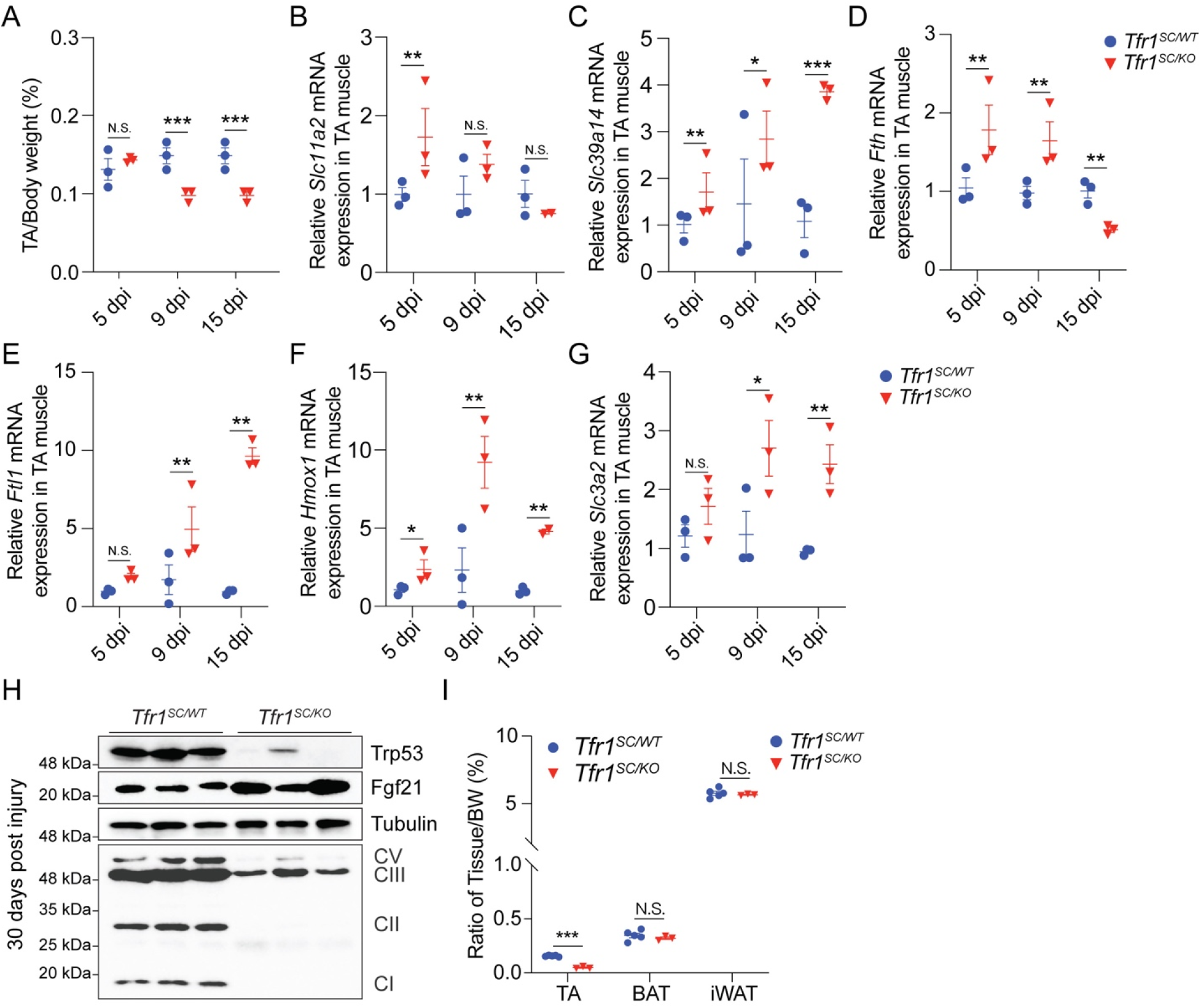
Related to Figure 6 Slc39a14-mediated NTBI absorption exacerbates skeletal muscle ferroptosis for *Tfr1^SC/KO^* mice. (A) Ratio of TA/body weight between *Tfr1^SC/WT^* and *Tfr1^SC/KO^* mice after CTX-induced injury at 5, 9 and 15 dpi; (B-G) qPCR analysis of *Slc11a2, Slc39a14, Fth1, Ftl, Hmox1, Slc3a2* expression in TA muscle of *Tfr1^SC/WT^* and *Tfr1^SC/KO^* mice after CTX-induced injury at 5, 9 and 15 dpi; (H) Representative western blot images of protein level in TA between *Tfr1^SC/WT^* and *Tfr1^SC/KO^* mice after CTX-induced injury at 30 dpi; (I) Ratio of tissue (TA, iBAT and iWAT)/body weight between *Tfr1^SC/WT^* and *Tfr1^SC/KO^* mice after CTX-induced injury at 30 dpi. N.S.: not significant, *P < 0.05, **P < 0.01, ***P < 0.005, by 2-sided Student’s t-test. Data represent the mean ± SEM.

**Figure S10.**
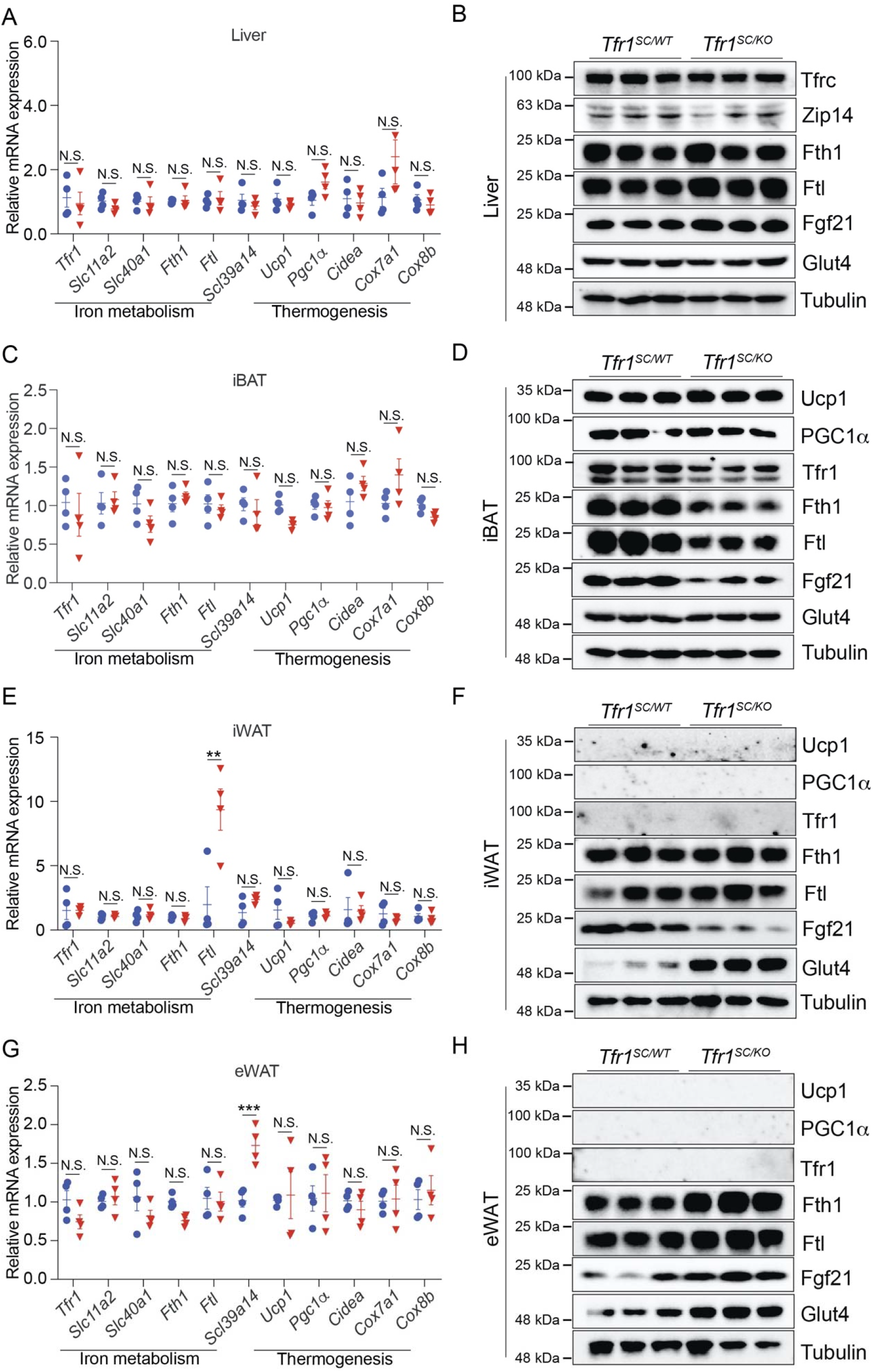
Related to Figure 6 SC-specific Tfr1 deletion induced TA muscle regeneration defect does not perturb systemic metabolism. (A, C, E and G) qPCR analysis of iron metabolism and thermogenesis related gene expression in Liver (A), iBAT (C), iWAT (E) and eWAT (G) between *Tfr1^SC/WT^* and *Tfr1^SC/KO^* mice after CTX-induced injury at 30 dpi. (B, D, F and. H) Representative western blot images of protein level in Liver (B), iBAT (D), iWAT (F) and eWAT (I) between *Tfr1^SC/WT^* and *Tfr1^SC/KO^* mice after CTX-induced injury at 30 dpi. N.S.: not significant, **P < 0.01, ***P < 0.005, by 2-sided Student’s t-test. Data represent the mean ± SEM (n=6/group).

## References and Notes

1. M. Raff, Cell suicide for beginners, Nature 396, 119–122 (1998).

2. C. Sciorati, E. Rigamonti, A. A. Manfredi, P. Rovere-Querini, Cell death, clearance and immunity in the skeletal muscle, Cell Death Differ. 23, 927–937 (2016).

3. Y. Tsujimoto, Multiple ways to die: non-apoptotic forms of cell *death*. Acta Oncol. 51, 293–300 (2012).

4. G. Corna, I. Caserta, A. Monno, P. Apostoli, A. A. Manfredi, C. Camaschella, P. Rovere-Querini, The Repair of Skeletal Muscle Requires Iron Recycling through Macrophage Ferroportin, J. Immunol. 197, 1914–1925 (2016).

5. S. J. Dixon, K. M. Lemberg, M. R. Lamprecht, R. Skouta, E. M. Zaitsev, C. E. Gleason, D. N. Patel, A. J. Bauer, A. M. Cantley, W. S. Yang, B. Morrison, B. R. Stockwell, Ferroptosis: An iron-dependent form of nonapoptotic cell death, Cell 149, 1060–1072 (2012).

6. W. S. Yang, B. R. Stockwell, Synthetic Lethal Screening Identifies Compounds Activating Iron-Dependent, Nonapoptotic Cell Death in Oncogenic-RAS-Harboring Cancer Cells, Chem. Biol. 15, 234–245 (2008).

7. Y. Xie, W. Hou, X. Song, Y. Yu, J. Huang, X. Sun, R. Kang, D. Tang, Ferroptosis: Process and function, Cell Death Differ. 23, 369–379 (2016).

8. N. Yagoda, M. Von Rechenberg, E. Zaganjor, A. J. Bauer, W. S. Yang, D. J. Fridman, A. J. Wolpaw, I. Smukste, J. M. Peltier, J. J. Boniface, R. Smith, S. L. Lessnick, S. Sahasrabudhe, B. R. Stockwell, RAS-RAF-MEK-dependent oxidative cell death involving voltage-dependent anion channels, Nature 447, 864–868 (2007).

9. X. Fang, H. Wang, D. Han, E. Xie, X. Yang, J. Wei, S. Gu, F. Gao, N. Zhu, X. Yin, Q. Cheng, P. Zhang, W. Dai, J. Chen, F. Yang, H. T. Yang, A. Linkermann, W. Gu, J. Min, F. Wang, Ferroptosis as a target for protection against cardiomyopathy, Proc. Natl. Acad. Sci. U. S. A. 116, 2672–2680 (2019).

10. F. Xuexian, C. Zhaoxian, W. Hao, H. Dan, C. Qi, Z. Pan, G. Feng, Y. Yingying, S. Zijun, W. Qian, A. Peng, H. Sicong, P. Jianwei, C. Hou-Zao, C. Jinghai, L. Andreas, M. Junxia, W. Fudi, Loss of Cardiac Ferritin H Facilitates Cardiomyopathy via Slc7a11-Mediated Ferroptosis, Circ. Res. 127, 486–501 (2020).

11. Y. Yu, L. Jiang, H. Wang, Z. Shen, Q. Cheng, P. Zhang, J. Wang, Q. Wu, X. Fang, L. Duan, S. Wang, K. Wang, P. An, T. Shao, R. T. Chung, S. Zheng, J. Min, F. Wang, Hepatic transferrin plays a role in systemic iron homeostasis and liver ferroptosis., Blood 136, 726–739 (2020).

12. H. Yin, F. Price, M. A. Rudnicki, Satellite cells and the muscle stem cell niche., Physiol. Rev. 93, 23–67 (2013).

13. H. Yin, F. Price, M. A. Rudnicki, Satellite Cells and the Muscle Stem Cell Niche, Physiol. Rev. 93, 23–67 (2013).

14. M. A. Hausburg, J. D. Doles, S. L. Clement, A. B. Cadwallader, M. N. Hall, P. J. Blackshear, J. Lykke-Andersen, B. B. Olwin, Post-transcriptional regulation of satellite cell quiescence by TTP-mediated mRNA decay, Elife 4 (2015), doi:10.7554/eLife.03390.

15. L. Xie, A. Yin, A. S. Nichenko, A. M. Beedle, J. A. Call, H. Yin, Transient HIF2A inhibition promotes satellite cell proliferation and muscle regeneration, J. Clin. Invest. 128, 2339–2355 (2018).

16. X. Yang, S. Yang, C. Wang, S. Kuang, The hypoxia-inducible factors HIF1alpha and HIF2alpha are dispensable for embryonic muscle development but essential for postnatal muscle regeneration., J. Biol. Chem. 292, 5981–5991 (2017).

17. C. Wang, W. Liu, Z. Liu, L. Chen, X. Liu, S. Kuang, Hypoxia Inhibits Myogenic Differentiation through p53 Protein-dependent Induction of Bhlhe40 Protein., J. Biol. Chem. 290, 29707–16 (2015).

18. T. Barrientos, I. Laothamatas, T. R. Koves, E. J. Soderblom, M. Bryan, M. A. Moseley, D. M. Muoio, N. C. Andrews, Metabolic Catastrophe in Mice Lacking Transferrin Receptor in Muscle., EBioMedicine 2, 1705–1717 (2015).

19. M. Altun, E. Edström, E. Spooner, A. Flores-Moralez, E. Bergman, P. Tollet-Egnell, G. Norstedt, B. M. Kessler, B. Ulfhake, Iron load and redox stress in skeletal muscle of aged rats, Muscle and Nerve 36, 223–233 (2007).

20. Y. Ikeda, A. Satoh, Y. Horinouchi, H. Hamano, H. Watanabe, M. Imao, M. Imanishi, Y. Zamami, K. Takechi, Y. Izawa-Ishizawa, L. Miyamoto, T. Hirayama, H. Nagasawa, K. Ishizawa, K. I. Aihara, K. Tsuchiya, T. Tamaki, Iron accumulation causes impaired myogenesis correlated with MAPK signaling pathway inhibition by oxidative stress, FASEB J. 33, 9551–9564 (2019).

21. R. Cui, S. E. Choi, T. H. Kim, H. J. Lee, S. J. Lee, Y. Kang, J. Y. Jeon, H. J. Kim, K. W. Lee, Iron overload by transferrin receptor protein 1 regulation plays an important role in palmitate-induced insulin resistance in human skeletal muscle cells, FASEB J. 33, 1771–1786 (2019).

22. M. Tieland, I. Trouwborst, B. C. Clark, Skeletal muscle performance and ageing., J. Cachexia. Sarcopenia Muscle 9, 3–19 (2018).

23. J. E. Levy, O. Jin, Y. Fujiwara, F. Kuo, N. C. Andrews, Transferrin receptor is necessary for development of erythrocytes and the nervous system., Nat. Genet. 21, 396–399 (1999).

24. L. Xie, A. Yin, A. S. Nichenko, A. M. Beedle, J. A. Call, H. Yin, Transient HIF2A inhibition promotes satellite cell proliferation and muscle regeneration, J. Clin. Invest. 128, 2339–2355 (2018).

25. T. Barrientos, I. Laothamatas, T. R. Koves, E. J. Soderblom, M. Bryan, M. A. Moseley, D. M. Muoio, N. C. Andrews, Metabolic Catastrophe in Mice Lacking Transferrin Receptor in Muscle., EBioMedicine 2, 1705–17 (2015).

26. S. Forsström, C. B. Jackson, C. J. Carroll, M. Kuronen, E. Pirinen, S. Pradhan, A. Marmyleva, M. Auranen, I. M. Kleine, N. A. Khan, A. Roivainen, P. Marjamäki, H. Liljenbäck, L. Wang, B. J. Battersby, U. Richter, V. Velagapudi, J. Nikkanen, L. Euro, A. Suomalainen, Fibroblast Growth Factor 21 Drives Dynamics of Local and Systemic Stress Responses in Mitochondrial Myopathy with mtDNA Deletions, Cell Metab. 30, 1040–1054.e7 (2019).

27. M. Ost, V. Coleman, A. Voigt, E. M. van Schothorst, S. Keipert, I. van der Stelt, S. Ringel, A. Graja, T. Ambrosi, A. P. Kipp, M. Jastroch, T. J. Schulz, J. Keijer, S. Klaus, Muscle mitochondrial stress adaptation operates independently of endogenous FGF21 action, Mol. Metab. 5, 79–90 (2016).

28. S. Welle, C. Thornton, R. Jozefowicz, M. Statt, Myofibrillar protein synthesis in young and old men., Am. J. Physiol. 264, E693–8 (1993).

29. A. M. Cuervo, J. F. Dice, Age-related decline in chaperone-mediated autophagy., J. Biol. Chem. 275, 31505–31513 (2000).

30. G. Shefer, D. P. Van de Mark, J. B. Richardson, Z. Yablonka-Reuveni, Satellite-cell pool size does matter: defining the myogenic potency of aging skeletal muscle., Dev. Biol. 294, 50–66 (2006).

31. C. S. Broome, A. C. Kayani, J. Palomero, W. H. Dillmann, R. Mestril, M. J. Jackson, A. McArdle, Effect of lifelong overexpression of HSP70 in skeletal muscle on age-related oxidative stress and adaptation after nondamaging contractile activity., FASEB J. Off. Publ. Fed. Am. Soc. Exp. Biol. 20, 1549–1551 (2006).

32. J. Palomero, A. Vasilaki, D. Pye, A. McArdle, M. J. Jackson, Aging increases the oxidation of dichlorohydrofluorescein in single isolated skeletal muscle fibers at rest, but not during contractions., Am. J. Physiol. Regul. Integr. Comp. Physiol. 305, R351–8 (2013).

33. S. H. Jung, L. R. DeRuisseau, A. N. Kavazis, K. C. DeRuisseau, Plantaris muscle of aged rats demonstrates iron accumulation and altered expression of iron regulation proteins, Exp. Physiol. 93, 407–414 (2008).

34. T. Hofer, E. Marzetti, J. Xu, A. Y. Seo, S. Gulec, M. D. Knutson, C. Leeuwenburgh, E. E. Dupont-Versteegden, Increased iron content and RNA oxidative damage in skeletal muscle with aging and disuse atrophy, Exp. Gerontol. 43, 563–570 (2008).

35. P. A. Leermakers, A. H. V. Remels, M. I. Zonneveld, K. M. A. Rouschop, A. M. W. J. Schols, H. R. Gosker, Iron deficiency-induced loss of skeletal muscle mitochondrial proteins and respiratory capacity; the role of mitophagy and secretion of mitochondria-containing vesicles, FASEB J. 34, 6703–6717 (2020).

36. M. Tkaczyszyn, M. Drozd, K. Węgrzynowska-Teodorczyk, I. Flinta, K. Kobak, W. Banasiak, P. Ponikowski, E. A. Jankowska, Depleted iron stores are associated with inspiratory muscle weakness independently of skeletal muscle mass in men with systolic chronic heart failure., J. Cachexia. Sarcopenia Muscle 9, 547–556 (2018).

37. A. Barberan-Garcia, D. A. Rodríguez, I. Blanco, J. Gea, Y. Torralba, A. Arbillaga-Etxarri, J. A. Barberà, J. Vilaró, J. Roca, M. Orozco-Levi, Non-anaemic iron deficiency impairs response to pulmonary rehabilitation in COPD., Respirology 20, 1089–1095 (2015).

38. J. F. Wilson, P. K. Joshi, Multivariate genomic scan implicates novel loci and haem metabolism in human ageing, Nat. Commun., 1–10 (2020).

39. J. Xu, M. D. Knutson, C. S. Carter, C. Leeuwenburgh, Iron accumulation with age, oxidative stress and functional decline, PLoS One 3 (2008), doi:10.1371/journal.pone.0002865.

40. K. C. DeRuisseau, Y.-M. Park, L. R. DeRuisseau, P. M. Cowley, C. H. Fazen, R. P. Doyle, Aging-related changes in the iron status of skeletal muscle, Exp. Gerontol. 48, 1294–1302 (2013).

41. C. Fillebeen, E. Charlebois, J. Wagner, A. Katsarou, J. Mui, H. Vali, D. Garcia-Santos, P. Ponka, J. Presley, K. Pantopoulos, Transferrin receptor 1 controls systemic iron homeostasis by fine-tuning hepcidin expression to hepatocellular iron load., Blood 133, 344–355 (2019).

42. J. Li, X. Pan, G. Pan, Z. Song, Y. He, S. Zhang, X. Ye, X. Yang, E. Xie, X. Wang, X. Mai, X. Yin, B. Tang, X. Shu, P. Chen, X. Dai, Y. Tian, L. Yao, M. Han, G. Xu, H. Zhang, J. Sun, H. Chen, F. Wang, J. Min, L. Xie, Transferrin Receptor 1 Regulates Thermogenic Capacity and Cell Fate in Brown / Beige Adipocytes, Adv. Sci. 1903366 (2020), doi:10.1002/advs.201903366.

43. S. Wang, X. He, Q. Wu, L. Jiang, L. Chen, Y. Yu, P. Zhang, X. Huang, J. Wang, Z. Ju, J. Min, F. Wang, Transferrin receptor 1-mediated iron uptake plays an essential role in hematopoiesis., Haematologica 105, 2071–2082 (2020).

44. J. Li, X. Pan, G. Pan, Z. Song, Y. He, S. Zhang, X. Ye, X. Yang, E. Xie, X. Wang, X. Mai, X. Yin, B. Tang, X. Shu, P. Chen, X. Dai, Y. Tian, L. Yao, M. Han, G. Xu, H. Zhang, J. Sun, H. Chen, F. Wang, J. Min, L. Xie, Transferrin Receptor 1 Regulates Thermogenic Capacity and Cell Fate in Brown/Beige Adipocytes, Adv. Sci. 7 (2020), doi:10.1002/advs.201903366.

45. C. Gaetano, L. Massimo, M. Alberto, Control of iron homeostasis as a key component of macrophage polarization, Haematologica 95, 1801–1803 (2010).

46. G. Corna, L. Campana, E. Pignatti, A. Castiglioni, E. Tagliafico, L. Bosurgi, A. Campanella, S. Brunelli, A. A. Manfredi, P. Apostoli, L. Silvestri, C. Camaschella, P. Rovere-Querini, Polarization dictates iron handling by inflammatory and alternatively activated macrophages, Haematologica 95, 1814–1822 (2010).

47. M. Marinkovic, C. Fuoco, F. Sacco, A. Cerquone Perpetuini, G. Giuliani, E. Micarelli, T. Pavlidou, L. L. Petrilli, A. Reggio, F. Riccio, F. Spada, S. Vumbaca, A. Zuccotti, L. Castagnoli, M. Mann, C. Gargioli, G. Cesareni, Fibro–adipogenic progenitors of dystrophic mice are insensitive to NOTCH regulation of adipogenesis, Life Sci. Alliance 2, e201900437 (2019).

48. A. Uezumi, S. I. Fukada, N. Yamamoto, S. Takeda, K. Tsuchida, Mesenchymal progenitors distinct from satellite cells contribute to ectopic fat cell formation in skeletal muscle, Nat. Cell Biol. 12, 143–152 (2010).

49. D. Kopinke, E. C. Roberson, J. F. Reiter, Ciliary Hedgehog Signaling Restricts Injury-Induced Adipogenesis, Cell 170, 340–351.e12 (2017).

50. X. Kang, M. Y. Yang, Y. X. Shi, M. M. Xie, M. Zhu, X. L. Zheng, C. K. Zhang, Z. L. Ge, X. T. Bian, J. T. Lv, Y. J. Wang, B. H. Zhou, K. L. Tang, Interleukin-15 facilitates muscle regeneration through modulation of fibro/adipogenic progenitors, Cell Commun. Signal. 16, 1–11 (2018).

51. J. E. Heredia, L. Mukundan, F. M. Chen, A. A. Mueller, R. C. Deo, R. M. Locksley, T. A. Rando, A. Chawla, Type 2 innate signals stimulate fibro/adipogenic progenitors to facilitate muscle regeneration, Cell 153, 376–388 (2013).

52. Y. Dong, K. A. S. Silva, Y. Dong, L. Zhang, Glucocorticoids increase adipocytes in muscle by affecting IL-4 regulated FAP activity, FASEB J. 28, 4123–4132 (2014).

53. B. Biferali, D. Proietti, C. Mozzetta, L. Madaro, Fibro-Adipogenic Progenitors Cross-Talk in Skeletal Muscle: The Social Network, Front. Physiol. 10, 1–10 (2019).

54. J. Huard, Y. Li, F. H. Fu, Muscle injuries and repair: current trends in research., J. Bone Joint Surg. Am. 84, 822–832 (2002).

55. A. J. De Micheli, E. J. Laurilliard, C. L. Heinke, H. Ravichandran, P. Fraczek, S. Soueid-Baumgarten, I. De Vlaminck, O. Elemento, B. D. Cosgrove, Single-Cell Analysis of the Muscle Stem Cell Hierarchy Identifies Heterotypic Communication Signals Involved in Skeletal Muscle Regeneration, Cell Rep. 30, 3583–3595.e5 (2020).

56. S. N. Oprescu, F. Yue, J. Qiu, L. F. Brito, S. Kuang, Temporal Dynamics and Heterogeneity of Cell Populations during Skeletal Muscle Regeneration, iScience 23, 100993 (2020).

57. M. Gao, P. Monian, Q. Pan, W. Zhang, J. Xiang, X. Jiang, Ferroptosis is an autophagic cell death process., Cell Res. 26, 1021–1032 (2016).

58. W. Hou, Y. Xie, X. Song, X. Sun, M. T. Lotze, H. J. 3rd Zeh, R. Kang, D. Tang, Autophagy promotes ferroptosis by degradation of ferritin., Autophagy 12, 1425–1428 (2016).

59. H. Feng, K. Schorpp, J. Jin, C. E. Yozwiak, B. G. Hoffstrom, A. M. Decker, P. Rajbhandari, M. E. Stokes, H. G. Bender, J. M. Csuka, P. S. Upadhyayula, P. Canoll, K. Uchida, R. K. Soni, K. Hadian, B. R. Stockwell, Transferrin Receptor Is a Specific Ferroptosis Marker, Cell Rep. 30, 3411–3423.e7 (2020).

60. A. W. B. Joe, L. Yi, A. Natarajan, F. Le Grand, L. So, J. Wang, M. A. Rudnicki, F. M. V. Rossi, Muscle injury activates resident fibro/adipogenic progenitors that facilitate myogenesis., Nat. Cell Biol. 12, 153–163 (2010).

61. A. A. Belaidi, A. I. Bush, Iron neurochemistry in Alzheimer’s disease and Parkinson’s disease: targets for therapeutics., J. Neurochem. 139 Suppl, 179–197 (2016).

62. M. Buijs, N. T. Doan, S. van Rooden, M. J. Versluis, B. van Lew, J. Milles, J. van der Grond, M. A. van Buchem, In vivo assessment of iron content of the cerebral cortex in healthy aging using 7-Tesla T2*-weighted phase imaging., Neurobiol. Aging 53, 20–26 (2017).

63. B. M. Owen, D. J. Mangelsdorf, S. A. Kliewer, Tissue-specific actions of the metabolic hormones FGF15/19 and FGF21, Trends Endocrinol. Metab. 26, 22–29 (2015).

64. L. J. Oost, M. Kustermann, A. Armani, B. Blaauw, V. Romanello, Fibroblast growth factor 21 controls mitophagy and muscle mass., J. Cachexia. Sarcopenia Muscle 10, 630–642 (2019).

65. A. Kharitonenkov, R. DiMarchi, FGF21 Revolutions: Recent Advances Illuminating FGF21 Biology and Medicinal Properties, Trends Endocrinol. Metab. 26, 608–617 (2015).

66. M. K. Badman, P. Pissios, A. R. Kennedy, G. Koukos, J. S. Flier, E. Maratos-Flier, Hepatic Fibroblast Growth Factor 21 Is Regulated by PPARα and Is a Key Mediator of Hepatic Lipid Metabolism in Ketotic States, Cell Metab. 5, 426–437 (2007).

67. T. Inagaki, P. Dutchak, G. Zhao, X. Ding, L. Gautron, V. Parameswara, Y. Li, R. Goetz, M. Mohammadi, V. Esser, J. K. Elmquist, R. D. Gerard, S. C. Burgess, R. E. Hammer, D. J. Mangelsdorf, S. A. Kliewer, Endocrine Regulation of the Fasting Response by PPARα-Mediated Induction of Fibroblast Growth Factor 21, Cell Metab. 5, 415–425 (2007).

68. F. F. Fisher, S. Kleiner, N. Douris, E. C. Fox, R. J. Mepani, F. Verdeguer, J. Wu, A. Kharitonenkov, J. S. Flier, E. Maratos-Flier, B. M. Spiegelman, FGF21 regulates PGC-1α and browning of white adipose tissues in adaptive thermogenesis, Genes Dev. 26, 271–281 (2012).

69. L. Kazak, E. T. Chouchani, M. P. Jedrychowski, S. P. Gygi, M. Bruce, L. Kazak, E. T. Chouchani, M. P. Jedrychowski, B. K. Erickson, K. Shinoda, P. Cohen, R. Vetrivelan, G. Z. Lu, D. Laznik-bogoslavski, S. C. Hasenfuss, S. Kajimura, S. P. Gygi, B. M. Spiegelman, Article A Creatine-Driven Substrate Cycle Enhances Energy Expenditure and Thermogenesis in Beige Fat Article A Creatine-Driven Substrate Cycle Enhances Energy Expenditure and Thermogenesis in Beige Fat, Cell 163, 643–655 (2015).

70. J. Z. Long, K. J. Svensson, L. A. Bateman, H. Lin, T. Kamenecka, I. A. Lokurkar, J. Lou, R. R. Rao, M. R. R. Chang, M. P. Jedrychowski, J. A. Paulo, S. P. Gygi, P. R. Griffin, D. K. Nomura, B. M. Spiegelman, The Secreted Enzyme PM20D1 Regulates Lipidated Amino Acid Uncouplers of Mitochondria, Cell 166, 424–435 (2016).

71. K. Ikeda, Q. Kang, T. Yoneshiro, J. P. Camporez, H. Maki, M. Homma, K. Shinoda, Y. Chen, X. Lu, P. Maretich, K. Tajima, K. M. Ajuwon, T. Soga, S. Kajimura, UCP1-independent signaling involving SERCA2b-mediated calcium cycling regulates beige fat thermogenesis and systemic glucose homeostasis., Nat. Med. 23, 1454–1465 (2017).

72. S. Keipert, D. Lutter, B. O. Schroeder, D. Brandt, M. Ståhlman, T. Schwarzmayr, E. Graf, H. Fuchs, M. H. De Angelis, M. H. Tschöp, J. Rozman, M. Jastroch, obesity resistance of UCP1-de fi cient mice, Nat. Commun. (2020), doi:10.1038/s41467-019-14069-2.

73. M. Z. Chen, J. C. Chang, J. Zavala-solorio, L. Kates, M. Thai, A. Ogasawara, X. Bai, S. Flanagan, V. Nunez, K. Phamluong, J. Ziai, R. Newman, FGF21 mimetic antibody stimulates UCP1-independent brown fat thermogenesis via FGFR1 / b Klotho complex in non-adipocytes, Mol. Metab. 6, 1454–1467 (2017).

74. L. M. Id, A. T. Id, M. De Bardi, F. F. C. Id, M. P. Id, G. R. D. Id, G. Imeneo, M. Bouch, L. Puri, F. D. S. Id, Macrophages fine tune satellite cell fate in dystrophic skeletal muscle of mdx mice, PLOS Biol., 1–29 (2019).

75. A. Mantovani, A. Sica, S. Sozzani, P. Allavena, A. Vecchi, M. Locati, The chemokine system in diverse forms of macrophage activation and polarization., Trends Immunol. 25, 677–686 (2004).

76. S. A. Villalta, H. X. Nguyen, B. Deng, T. Gotoh, J. G. Tidball, Shifts in macrophage phenotypes and macrophage competition for arginine metabolism affect the severity of muscle pathology in muscular dystrophy., Hum. Mol. Genet. 18, 482–496 (2009).

77. B. Vidal, A. L. Serrano, M. Tjwa, M. Suelves, E. Ardite, R. De Mori, B. Baeza-Raja, M. Martínez de Lagrán, P. Lafuste, V. Ruiz-Bonilla, M. Jardí, R. Gherardi, C. Christov, M. Dierssen, P. Carmeliet, J. L. Degen, M. Dewerchin, P. Muñoz-Cánoves, Fibrinogen drives dystrophic muscle fibrosis via a TGFbeta/alternative macrophage activation pathway., Genes Dev. 22, 1747–1752 (2008).

78. A. C. Thomas, W. J. Eijgelaar, M. J. A. P. Daemen, A. C. Newby, The pro-fibrotic and anti-inflammatory foam cell macrophage paradox, Genomics data 6, 136–138 (2015).

79. D. Kim, B. Langmead, S. L. Salzberg, HISAT: A fast spliced aligner with low memory requirements, Nat. Methods 12, 357–360 (2015).

80. S. Anders, P. T. Pyl, W. Huber, HTSeq—a Python framework to work with high-throughput sequencing data, Bioinformatics 31, 166–169 (2015).

81. M. I. Love, W. Huber, S. Anders, Moderated estimation of fold change and dispersion for RNA-seq data with DESeq2, Genome Biol. 15, 1–21 (2014).

82. G. Yu, L.-G. Wang, Y. Han, Q.-Y. He, clusterProfiler: an R Package for Comparing Biological Themes Among Gene Clusters, Omi. A J. Integr. Biol. 16, 284–287 (2012).

83. A. Subramanian, P. Tamayo, V. K. Mootha, S. Mukherjee, B. L. Ebert, M. A. Gillette, A. Paulovich, S. L. Pomeroy, T. R. Golub, E. S. Lander, J. P. Mesirov, Gene set enrichment analysis: A knowledge-based approach for interpreting genome-wide expression profiles, Proc. Natl. Acad. Sci. 102, 15545 LP – 15550 (2005).

84. N. Das, L. Xie, S. K. Ramakrishnan, A. Campbell, S. Rivella, Y. M. Shah, Intestine-specific Disruption of Hypoxia-inducible Factor (HIF)-2alpha Improves Anemia in Sickle Cell Disease., J. Biol. Chem. 290, 23523–23527 (2015).

85. X. Pan, B. Liu, S. Chen, H. Ding, X. Yao, Y. Cheng, D. Xu, Y. Yin, X. Dai, J. Sun, G. Xu, J. Pan, L. Xiao, L. Xie, Nr4a1 as a myogenic factor is upregulated in satellite cells/myoblast under proliferation and differentiation state., Biochem. Biophys. Res. Commun. 513, 573–581 (2019).

86. S. Chen, H. Ding, X. Yao, L. Xie, Isolation and Culture of Single Myofiber and Immunostaining of Satellite Cells from Adult C57BL/6J Mice, Bio-protocol 9, e3313 (2019).

